# Molecular and Spatial Organization of the Primary Olfactory System and its Responses to Social Odors

**DOI:** 10.1101/2025.05.02.651832

**Authors:** Bogdan Bintu, Yoh Isogai, Xiaowei Zhuang, Catherine Dulac

## Abstract

The detection of olfactory cues is essential to signal food, predators, and social encounters. To determine how the sensory detection of physiologically relevant odors is systematically mapped into the mouse primary olfactory system, we used Multiplexed Error Robust Fluorescent *In Situ* Hybridization (MERFISH) to construct a molecular atlas of odorant receptor (OR) expression in the main olfactory epithelium (MOE) and olfactory bulb (OB). We comprehensively quantified the expression of the mouse OR repertoire and uncovered stereotypical gradients of sensory neuron distribution in the MOE along two, central-to-peripheral and basal-to-apical, axes. Projections of sensory neurons mirror MOE gradients along the dorsal-ventral and anterior-posterior axes of the OB, respectively. Integration with sequencing data revealed candidate signaling molecules underlying this spatial organization. Co-imaging OR and activity marker expression identified distinct spatial domains of sensory responses in the MOE and OB, providing a topographical basis for olfactory responses to ethologically relevant odors.

## Introduction

In mammals, the main olfactory epithelium (MOE) detects a vast range of odor signals conveying information about the environment, such as the presence of food, predators, and conspecifics^1,2^. Within the MOE, odor cues are detected by olfactory sensory neurons (OSNs) in which an intricate mode of transcriptional regulation leads to the expression of a single olfactory receptor (ORs) gene out of a large ∼1000 OR gene family in rodents^3,4^. Commitment to a given OR expression is orchestrated by coordinated transcriptional regulation and chromatin modifications^5–8^. Expression of specific combinations of axonal guidance and signaling molecules enables olfactory neurons expressing a given OR to send precise axonal projections into distinct pairs of glomeruli in the olfactory bulb (OB), generating an organized map of odor sensing ^9–11^.

Past efforts have revealed some of the rules governing the spatial organization of OSNs within the MOE and their axonal projections to the OB. In the MOE, expression of individual OR genes appears restricted within concentric olfactory zones whose position is highly stereotypical across individuals in a given species^12,13^. While 4-5 discrete olfactory zones were initially defined based on expression of a few ORs *in situ*^12,13^, additional *in situ* measurements uncovered more, partially overlapping olfactory zones^14,15^. In turn, spatial transcriptomics approaches have begun to provide a more comprehensive, yet so far low spatial resolution view of OR organization within the MOE^16,17^ and suggested a more continuous, rather than discrete spatial distribution of distinct ORs along the central-peripheral axis of the MOE. The OR distribution along this axis is associated with gradients of specific transcription factors, which, when genetically perturbed, were shown to alter the organization of OR expression in the MOE^18^.

Within the OB, specific OR transcripts are localized to sensory axon terminals in bilaterally symmetric pairs of glomeruli – one medial and one lateral per hemisphere^9,10^. The highly stereotyped positions of specific OR glomeruli across animals suggested the existence of a chemotopic map of OR activation in the OB^1,2,9,10,19^. These findings were confirmed with a larger set of ORs using transgenic strategies^11,20,21^ as well as newly developed spatial transcriptomics approaches ^22,23^. However, these OR mapping efforts either encompassed less than 10% of the OR repertoire due to low *OR* detection efficiency or relied on computational reconstructions with low spatial resolution due to averaging measurements across multiple animals. Thus, a rigorous and systematic understanding of the precise OR representation and its stereotypy within the glomeruli layer of the OB remains to be achieved.

The molecular stereotypy of sensory neuron projection at fixed positions of the OB uncovered a spatial map underlying odor sensing^1,2,9,10^ and functional imaging of the dorsal side of the OB in animals exposed to discrete chemical compounds revealed distinct topographical domains localized in sub-regions of the OB^24–29^. However, progress in connecting the spatial organization of OSNs or their axonal projections to their sensory function is impeded by the lack of complete molecular atlases detailing the OR expression in the MOE and OB and the lack of functional annotations of each OR to physiologically relevant cues. While many methodologies have been developed to catalogue OR responses to different olfactory cues^30^, including recent, high throughput, *in vivo* screens^31–34^, these methods typically characterized OR responses to unnaturally high concentrations (mM range) of single compounds and have yet to comprehensively describe the OR responses to naturalistic and physiologically relevant odors normally encountered by an animal.

In this study we took advantage of the sensitivity of the spatial transcriptomics method Multiplexed Error Robust Fluorescent *In Situ* Hybridization (MERFISH)^35^ to assess the expression of the near-complete ∼1100 OR repertoire and ∼10 TAAR receptors and construct detailed spatial atlases of the receptor distribution within the MOE and the OB. By further combining MERFISH with single molecule FISH for *Egr1*, a marker of neuronal activity in chemosensory organs^36,37^ we characterized the genome-wide response of ORs to ethologically-relevant odorants, with a focus on social odors. The integration of the spatial organization of ORs with their functional characteristics uncovered the existence of spatial domains responding to distinct physiologically relevant olfactory signals.

## Results

### *In situ* mapping of the near-complete repertoire of odorant receptor gene expression in the MOE using MERFISH

MERFISH is a single-cell transcriptome imaging approach that allows localization and quantification of RNAs in single cells, in situ, at the scale of hundreds to thousands of transcripts^35,38,39^. To detect the large repertoire of ∼1000 odorant receptors, two modifications were added to the standard MERFISH protocol^38^. First, to overcome the high sequence similarity between the coding sequences of OR and Taar genes, MERFISH probes (Table S1) were designed to preferentially target the more variable untranslated regions (UTRs) whose annotations were recently refined^40^ (Figure S1 A,B). Second, to increase detection sensitivity, we employed a combination of split probes^41^ and multiplexed branched DNA amplification^42^ (Figure S1 C).

For the MOE of each mouse (N=8 males and N=5 females) six coronal positions along the anterior-posterior axis were selected. For each anterior-posterior position, 2-5 consecutive 16-µm sections were imaged via MERFISH at the scale of 200-500 ORs per section (Figure 1 A-C). These consecutive sections were aligned to each other in order to detect the expression of the entire ∼1000 OR/Taar gene repertoire.

**Figure 1.**
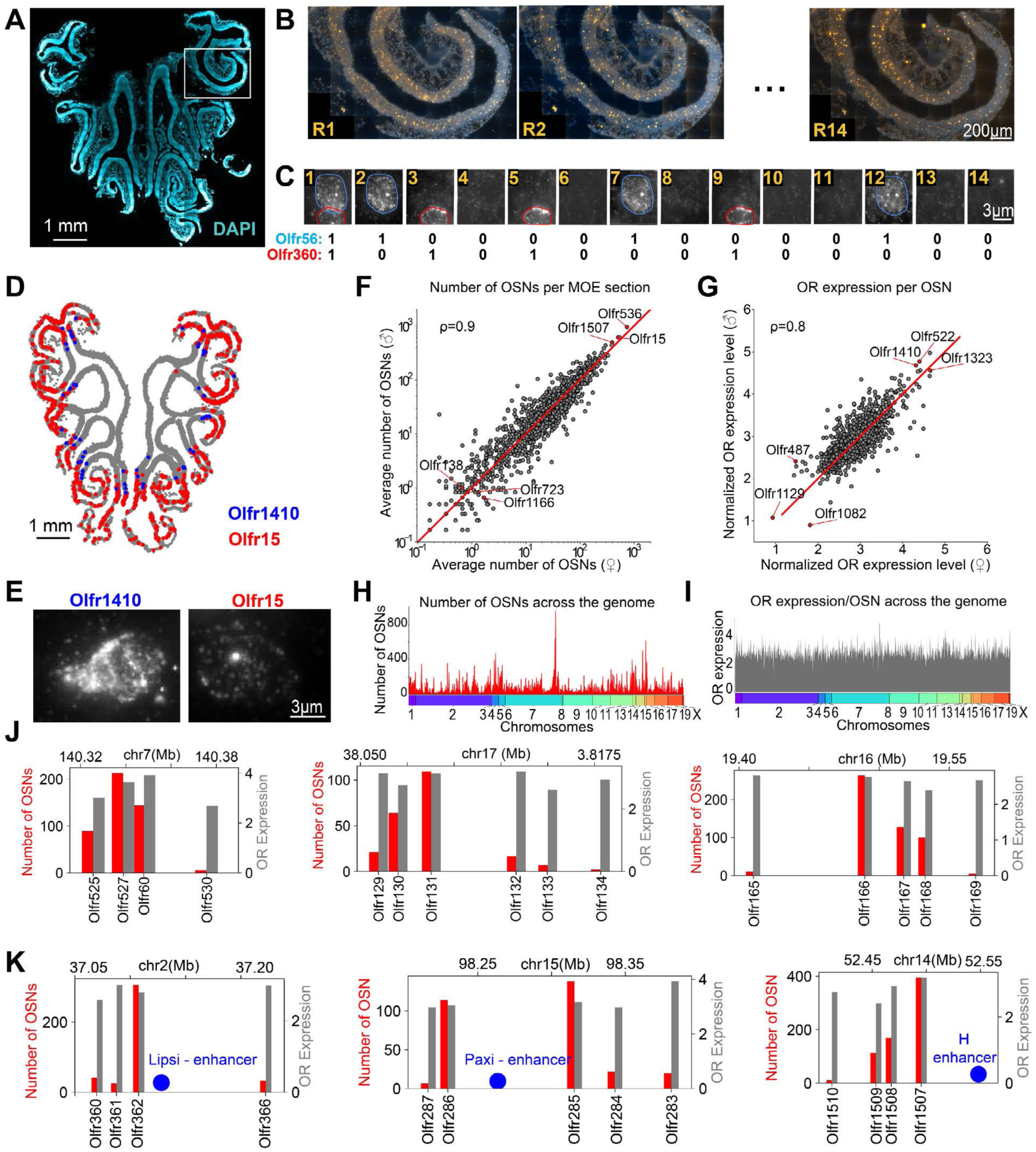
MERFISH enables high-throughput imaging of olfactory sensory neurons in the MOE at the OR-repertoire scale. **(A)** Image of a 15-µm coronal section of the MOE with DAPI stained nuclei (blue). Encoding MERFISH probes designed for a set of 200 OR genes were hybridized and detected by sequential rounds of imaging with readout probes complementary to the readout sequences on encoding probes, allowing decoding of ORs. **(B)** Images of a subregion of the section in (A) showing the fluorescent signal (yellow) of individual MERFISH readouts. **(C)** Images of the 14 readout MERFISH cycles for a small subregion of the MOE containing two sensory neurons identified as Olfr56 and Olfr360 based on the unique combination of readout detection. **(D)** Coronal MOE section (gray) overlayed with the positions of two types of OSNs expressing Olfr1410 (blue) or Olfr15 (red). **(E)** Representative MERFISH images for an Olfr1410 OSN and an Olfr15 OSN. **(F)** Correlation between the average number of OSNs per section identified for each OR in female vs male animals. **(G)** Correlation between the average expression level per OSN for each OR for female vs male animals. The expression level was estimated from the brightness of the MERFISH fluorescent signal and normalized by the average brightness across the entire MOE **(H)** Average number of OSNs / section and **(I)** Average expression level per OSN for each OR ordered by the genomic position of each OR. The bottom color label indicates the corresponding chromosome. **(J)** Bar plots indicating the number of OSNs per slice (red) and the OR expression level per OSN (gray) for representative OR genomic clusters. **(K)** Same as (J) for OR genomic clusters close to known enhancer elements (blue).

To estimate the efficiency and accuracy of MERFISH detection for specific receptor genes, we performed control experiments using MOE sections from two transgenic lines Olfr17-IRES-tau-lacZ^11^ (N=1) and Olfr16-IRES-tau-GFP^43^ (N=1) in which neurons expressing the OR genes Olfr17 or Olfr16 co-expressed the reporter genes beta-galactosidase (β-gal) or green fluorescent protein (GFP), respectively. We analyzed two sections per animal and found that 75 and 77% of cells expressing β-gal/GFP were identified by MERFISH as Olfr17 and Olfr16 positive, respectively, and that 95 and 98% of cells identified as expressing Olfr17 and Olfr16, respectively, by MERFISH also expressed β-gal/GFP (Figure S2). For the hundreds of receptors targeted in each MOE section, these results indicate high efficiency and accuracy of our experimental paradigm in quantifying the OR identity of individual OSNs.

### Representation and expression levels of ORs in the MOE

Based on the acquired MERFISH data, the average number of OSNs per section expressing a given receptor and the average receptor expression levels within the corresponding OSN population were quantified across ∼200 coronal sections sampling the anterior-posterior axis of the MOE across 13 animals (N=8 male and N=5 female animals) (Figure 1D-G and Table S2). While the absolute number of OSNs varied between anterior, central, or posterior MOE sections, the relative proportions of OSN types and the average level of OR expression per cell remained similar along the antero-posterior axis of the MOE (Figure S3). The number of OSNs per section and receptor expression level per cell were also similar between males and females across both ORs and Taars (Pearson correlation coefficients of 0.9 and 0.8, respectively, Figure 1 F,G). These data are consistent with analysis of OR expression in female and male mice by bulk-RNA sequencing^40^. By contrast, large variations were found in the number of OSNs expressing a given OR across the OR repertoire, spanning three orders of magnitude from the most to the least abundant OR populations (Figure 1F). For instance, the most abundant 5 ORs (including Olfr536, Olfr15 and Olfr1507) were expressed in >350 OSNs per MOE section on average while 5.2% of receptors (including Olfr1166, Olfr138 or Olfr723) are sparsely expressed and detected in <1 OSNs per slice on average. By contrast, expression levels of ORs per cell were more uniform, varying by less than an order of magnitude from the lowest expressed ORs (i.e. Olfr1129, Olfr487, or Olfr1082) to the most expressed ORs (Olfr1323, Olfr522, or Olfr1410), with most ORs exhibiting expression levels within a factor of two of each other (Figure 1G).

Next, we assessed the various neuronal populations according to the genomic coordinates of their expressed OR. *OR*s appear clustered within the genome with OSN numbers forming peaks and valleys: OSN population sizes progressively decreased with the genomic distance from an abundant OSN type (Figure 1H-J, Figure S3J). By contrast, the expression levels per cell of the corresponding ORs had only a weak correlation with the OSN population sizes (Pearson correlation of 0.1) and were less variable across the genome (Figure 1H-J, Figure S3K). Notably, ORs located within 50 kb of known super-enhancers (i.e. the *H-element* or the *Greek islands enhancers*)^44^ (Figure 1 K) were expressed by higher numbers of sensory neurons than other ORs (p-value 8.3×10^−5^, Wilcoxon test) consistent with an enhancer-based regulation of OR choice.

### Spatial organization of the OR repertoire within the MOE

Analysis of the spatial distribution of OR expression within the MOE showed that, as previously described, certain ORs are expressed in closer spatial proximity to each other and within distinct subregions of the MOE (Figure 2A). To quantify this phenomenon, we systematically calculated the degree of spatial overlap for each pair of ORs across 13 animals. The degree of overlap between two OSN types was defined as the fraction of cells located within a cutoff distance of 200µm, averaged across all the MOE sections imaged. Each OSN type was then plotted in a 2D UMAP where OSNs with higher degree of spatial overlap are mapped closer to each other. Within the UMAP space, ORs defined a thin curved distribution (Figure 2B, Table S3). By ordering the ORs according to this UMAP coordinate, we noticed that each OSN type overlapped with multiple adjacent OSN types in a graded fashion (Figure 2C) suggestive of a continuous distribution of OR expression within the MOE.

**Figure 2.**
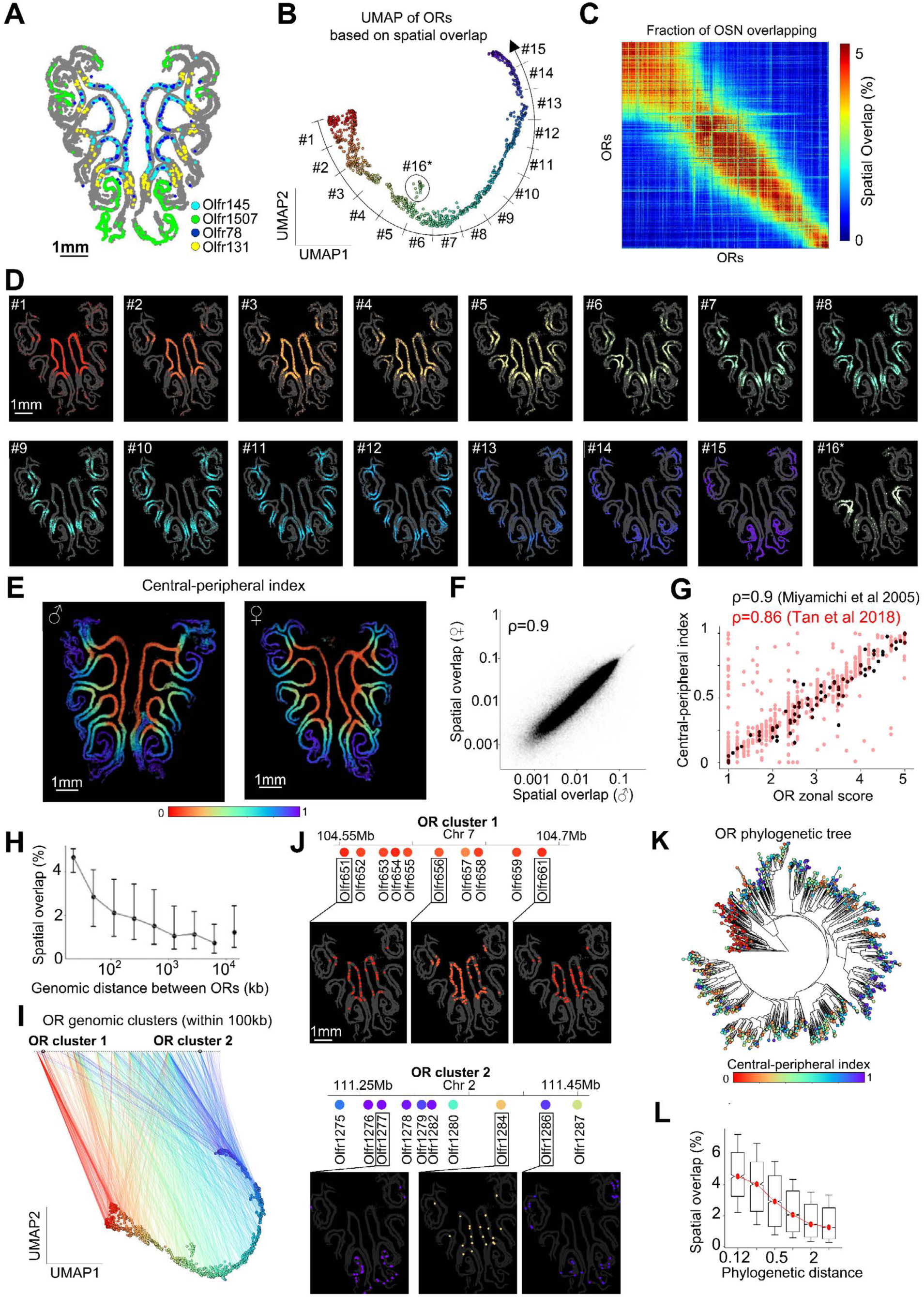
The spatial distribution of sensory neurons in the MOE. **(A)** Image of a central coronal section of the MOE of a male mouse (in gray) showing the position of four example OSN types (Olfr145, Olfr1507, Olfr78, Olfr131) marked in different colors.. **(B)** UMAP embedding of OSN types based on spatial overlap. Smaller distances between OSNs reflect higher degree of spatial overlap in the MOE. ORs are colored according to their position along the curved UMAP curved trajectory. The normalized position (ranging 0-1) along this trajectory for each OR is defined as the central-peripheral index. **(C)** Matrix quantifying the degree of spatial overlap in the MOE between different types of OSNs. Spatial overlap was defined pairwise for OSNs types as the fraction of pairs of cells within a cutoff distance of 200 µm. The ORs were ordered based on the central-peripheral index from (B). **(D)** Image of an MOE section onto which groups of ∼70 ORs progressively covering the UMAP coordinate in (B) are shown. The last image (#16 marked with *) covers the unusual zone of ORs. **(E)** Two examples of MOE sections (from a male and female) with the OSNs colored based on the central-peripheral index from (B). **(F)** Correlation between spatial overlap of OSN types in females and males. **(G)** Correlation between the central-peripheral index measured by MERFISH and the continuous zonal score measured across previous studies^15,16^ **(H)** Correlation between spatial overlap and genomic distance of ORs. **(I)** Line plot connecting each OR on the UMAP (B) to its corresponding genomic OR cluster. ORs were clustered across the genome if located within 100kb of each other. **(J)** Spatial positions of OSNs across two examples of genomic clusters. **(K)** Phylogenetic tree connecting ORs based on their sequence similarity (constructed as in ^32^). Each node, representing an OR, is colored based on the central-peripheral index from (B). **(L)** Correlation between the distance along the phylogenetic tree of pairs of ORs and their spatial overlap. Notches represent 95% confidence intervals; the box marks the first and third quartile; whiskers mark the 15th and the 85th percentiles.

In a different representation, each OSN was displayed within MOE sections with a color scheme reflecting the UMAP coordinate of the corresponding OR type. This representation revealed the main mode of organization of ORs as overlapping rings that continuously expand from the center to the periphery (Figure 2D). This organization was highly stereotypical across animals, including males and females (Figure 2E, F). While consistent with a recent report^14^ showing the expression of each OR constrained to ring-like zones with different rings having varying degrees of overlap, our near complete screening of the OR repertoire reveals that the positions of these rings change in a continuous fashion, impeding the definition of precise zonal boundaries (Figure 2F). A noticeable exception to this rule was found for a small group of ORs displaying a separate distribution from the main UMAP coordinate and, rather than forming a ring, were constrained to a mid-region of the MOE (Figure 2B, D marked with *) previously characterized as a separate, unusual zone^45^. In contrast to ORs, Taars did not form a continuous spatial distribution but rather were biased towards the central region of the MOE or to a mid-peripheral ring (Figure S4A, B).

Validating these results, a close correspondence was observed between the distribution of ORs quantified by MERFISH along the central-peripheral axis of the MOE (0-denoting most central and 1 most peripheral location) and the published zonal annotations (1-5), based on non-multiplexed *in situ* hybridizations of ∼80 ORs^15^ (Figure 2G – black dots, Pearson correlation of 0.9). Similarly, a high correspondence (Pearson correlation of 0.86) was found between our MERFISH results across the OR repertoire and a sequencing-based approach of inferring the OR zonal annotation^16^ (Figure 2G – red dots).

We next explored the relationship between the spatial organization of different *ORs* in the MOE and their position within the genome and sequence similarity based on a previously published phylogenetic tree^32^. Our analysis showed that *ORs* located in proximity to each other within the genome (i.e., within 100kb) had higher spatial overlap than more distal ORs (Figure 2H). Accordingly, the expression of ORs belonging to the same genomic cluster was predominantly constrained within the same ring-like structure in the MOE (Figure 2I-K). For instance, the cluster of receptors Olfr651-Olfr661 were all expressed within the central ring of the MOE. However, we also noted that many OR genomic clusters had a few OR exceptions which were not expressed within the same MOE region as the rest of the cluster. For instance, within cluster Olr1275-Olfr1287, most ORs were constrained to the same peripheral ring in the MOE, but Olfr1284 and Olfr1287 were exceptions, with more central expression patterns (Figure 2J).

Additionally, ORs sharing high sequence similarity, and hence are close phylogenetically (Figure 2K, L), had higher overlap and were predominantly expressed within the same MOE ring structures. These results are consistent with the view that spatially patterned transcription factors^18^ recruited to similar sequences in the OR gene body and/or shared neighboring regulatory sequences enable ORs to be coregulated spatially. Further mapping of transcription factor binding sites and enhancer-promoter interactions^7,18,44^ may help establish a comprehensive model of how OR expression is spatially regulated.

### Mapping the projections of OR sensory neurons to the olfactory bulb

We took advantage of the high sensitivity of MERFISH to systematically track OR transcripts in OSN axon terminals within olfactory bulb glomeruli. MERFISH was performed on serially sectioned bilateral olfactory bulbs from 2 adult females (Figure 3A). Glomeruli were segmented in each 2D section based on the nuclear signal from surrounding periglomerular cells using neural network-based models^46^. The expression of single *ORs* with higher than 10 detected transcripts was identified in only 1-3 glomeruli per hemi-bulb spanning a few OB sections (Figure 3B, C). Glomerular OR identity was determined based on the most expressed OR in each glomerulus with >10 detected transcripts. Data obtained from serial sections of bilateral bulbs were aligned and stacked together to reconstruct a 3D projection map of OSNs to the OB across the OR and Taar repertoire (Figure 3D, Figure S4C). For each OB imaged ∼2000 glomeruli were identified which covered collectively 641 receptors.

**Figure 3.**
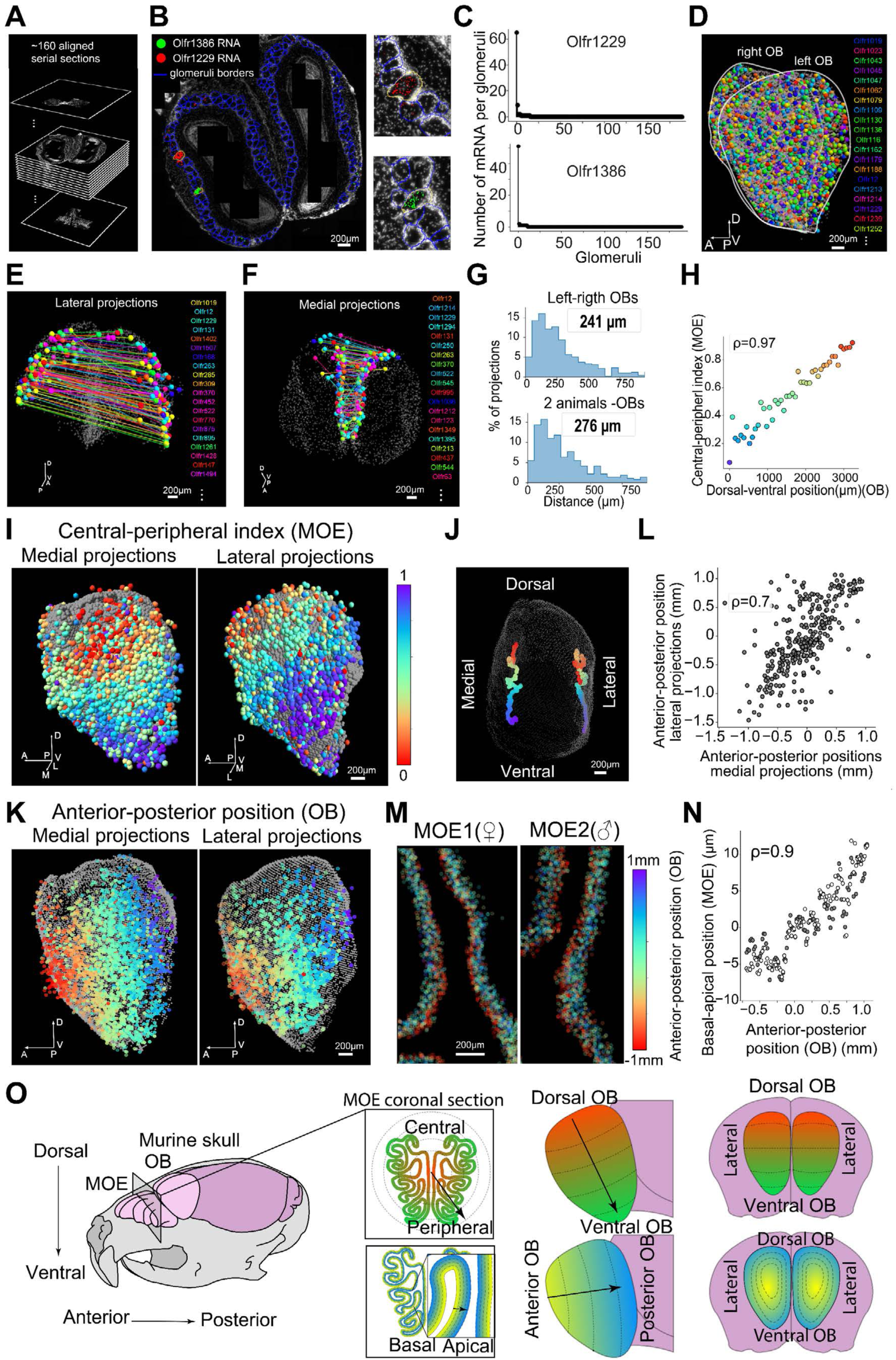
3D projection of OSNs into the OB. **(A)** A set of ∼160 serial coronal sections (16-18µm thick) covering almost continuously the bilateral OBs of one mouse were imaged with MERFISH using OR probes. DAPI nuclear staining was used to align the sections into a 3D stack. **(B)** Image of one of the mid-sections in the 3D stack in (A). Blue lines mark the boundaries of glomeruli segmented using a network-based machine learning approach based on the nuclear signal of periglomerular cells (white)^46^. Transcripts identified by MERFISH for two example *ORs*, *Olfr1386* (green) and *Olfr1229* (red), were overlaid. Glomeruli enriched in each OR are enlarged on the right. **(C)** Distribution of the number of transcripts for Olfr1386 (top) and Olfr1229 (bottom) across the glomeruli segmented in the section shown in (B). **(D)** 3D map of the OB reconstructed in (A). Glomeruli are displayed as spheres colored based on OR identity. **(E)** 3D map in (D) with only laterally projecting ORs highlighted. Lines connect projections with the same OR-identity between left and right bilateral bulbs. **(F)** Same as in (E) for medial projections. **(G)** Histograms of distances between glomeruli of matching OR identity across bilateral bulbs (top) or bulb of different animals (bottom). The median inter-glomeruli distances were highlighted for each histogram. **(H)** Correlation between the central-peripheral index of ORs in the MOE and the dorsal-ventral position of OR projections into the OB. OB projections were binned across 150µm intervals along the dorsal-ventral axis. **(I)** Consensus 3D map with aligned OR projections across 4 imaged OBs. Projections are colored based on the central-peripheral index of their corresponding OR as defined in Figure 2B. The lateral and medial projections are displayed on the left and right respectively. **(J)** 3D map of the OB (gray dots) onto which we marked the center positions of projections across groups ∼70 ORs. Each group is defined by a sliding window of size 70 across the ORs ordered according to the central-peripheral index in the MOE. **(K)** Left: Medial OB projections colored according to their anterior-posterior position in the OB. Right: Lateral OB projections colored according to medial anterior-posterior position. **(L)** Correlation across ORs of the anterior-posterior positions between their medial and lateral projections. **(M)** MOE sections of a male (left) and a female mouse (right) with OSNs colored according to the anterior-posterior position of their corresponding OB projection (defined in the left panel of (K)). **(N)** Correlation across OSN types between the average anterior-posterior position of their projections into the OB and the average basal-apical position in the MOE. The quantifications across MOE sections from a male and from a female are shown separately in gray and white respectively. **(O)** Schematic summarizing the two axes of organization of OSNs in the MOE and their projections in the OB.

To validate the accuracy of the OB projection map, we selected 5 ORs (Olfr15, Olfr16, Olfr17, Olfr155, and Olfr1507) whose 3D projections were previously determined using transgenic reporters^20^ (Figure S5A). The comparison of pairwise distances between the OB projections of these ORs in our MERFISH dataset and the published report demonstrated a tight correspondence (Figure S5A, B, C, Pearson correlation coefficient of 0.94).

Comparison of the OB projections from the left and right hemispheres of individual animals revealed a striking mirror-symmetry for both medial and lateral projections (Figure 3E, F). Quantitatively, upon aligning left and right OBs, the 3D positions of the projections for corresponding ORs matched within ∼250 µm (Figure 3G – top panel). Similar results were obtained when comparing the bulbs imaged from 2 animals (median distance 276 µm), indicating that the projections are spatially specified across the OB within an accuracy of ∼3 glomeruli diameters on average across the OR repertoire (Figure 3G – bottom panel). Based on this stereotypy, a consensus projection map was constructed by aligning the four OBs imaged into a common reference. The alignment enabled a more relaxed OR identification criterion, permitting glomeruli with >5 transcripts to be assigned to an OR if the same OR identities were observed in a similar position across multiple OBs (<500 µm). This approach increased the coverage to approximately 75% of the OR repertoire (Table S4).

### Spatial organization of ORs in the MOE versus projections to the OB

We next asked how the spatial distributions of ORs in the MOE and their projections to the OB relate to each other. Mapping OR projections into the OB according to the central-peripheral index of OR expression in the MOE (used in Figure 2B, D, E) revealed that the continuous rings in the MOE mapped to continuous bands in the OB along the dorsal to ventral direction (Figure 3H). A tight correlation was found between the dorsal-ventral location in the OB and the central-peripheral index in the MOE of corresponding ORs (Pearson correlation coefficient of 0.97) (Figure 3I). Likewise, averaging the 3D position of projections along a sliding window of ∼80 ORs along the central-peripheral MOE direction revealed the emergence of two parallel dorsal-ventral axes (one for medial and one for lateral projections) in the OB. These results suggest that the central-peripheral preference of ORs in the MOE is transferred to a dorsal-ventral preference of OR projections into the OB, with symmetry across the medial and lateral sides of the OB.

We then examined the distribution of OR projections along the orthogonal anterior-posterior axis of the olfactory bulb. As seen with the dorsal-ventral axis, the anterior-posterior distribution of OR projections was mirrored between medial and lateral glomeruli (Figure 3K, L Pearson correlation coefficient of 0.8). Surprisingly, when mapping this anterior-posterior distribution of OR projections from the OB into OSNs in the MOE, we noticed an unexpected basal-to-apical distribution of OR expression in the MOE neuroepithelium (Figure 3M, N) with some ORs (i.e. Olfr1170 and Olfr1388) expressed consistently closer to the lumen (apical) while others (i.e. Olfr354 and Olfr99) were expressed closer to the basal lamina (basal) (Figure S6A-I). This distribution (Table S5) was conserved for both central and peripheral ORs across anterior, central and posterior sections of the MOE (Figure S6A-I) Pearson correlation coefficient 0.8). When mapping the MOE basal-to-apical preference of ORs onto the OB we noticed the emergence of two symmetrical axes (one for medial and one for lateral glomeruli) aligned with the anterior-posterior axis of the OB (Figure S6J-L). Finally, and consistent with observations in the MOE (Figure 2), ORs more similar in sequence or closer in genomic space were associated with closer OB projections along both the dorsal-ventral and anterior-posterior axes of the OB (Figure S7A-D).

Together, these results uncover two sets of preferential axes of OR distribution in both the MOE and OB (Figure 3O). In the MOE, in addition to the expected large scale central-peripheral preference (Figure 2 B-D), the distribution of ORs also displayed a smaller scale basal-apical preference. These two gradients of OSNs in the MOE were mirrored to two corresponding axes (dorsal-ventral and anterior-posterior) of distributions of OSN projections in the OB, where the OSNs with a more central or basal preference in the MOE projected to the more dorsal or anterior regions of the OB, respectively.

### Differential gene expression in OSNs according to OR spatial distribution in the MOE and OB

The spatial distribution of ORs within the MOE and OB results from the coordinated expression of multiple genes encoding axonal guidance molecules, transcription factors and other signaling molecules^1,47^. To correlate the differential gene expression in OSNs expressing distinct receptors with their spatial distribution in the MOE and their OB projections, we integrated published single-cell RNA sequencing data from adult OSNs^47^ with MOE and OB spatial maps generated by MERFISH (Figure 4A). For a given gene, the average scRNAseq expression level across OSNs of each OR type was imputed onto the MOE location of OSNs or onto the OB projections of the corresponding OR type identified by MERFISH (Figure 4A, B and Methods). These imputations correctly approximated the expression patterns previously measured for individual genes in the MOE or the OB. For instance, the cell adhesion molecule *Ncam2* and the quinone reducing enzyme *Nqo1*, both in our imputations (Figure 4B) and in previous immunofluorescence images^48^, showed complementary central-peripheral and dorsal-ventral distributions in the MOE and OB, respectively.

**Figure 4.**
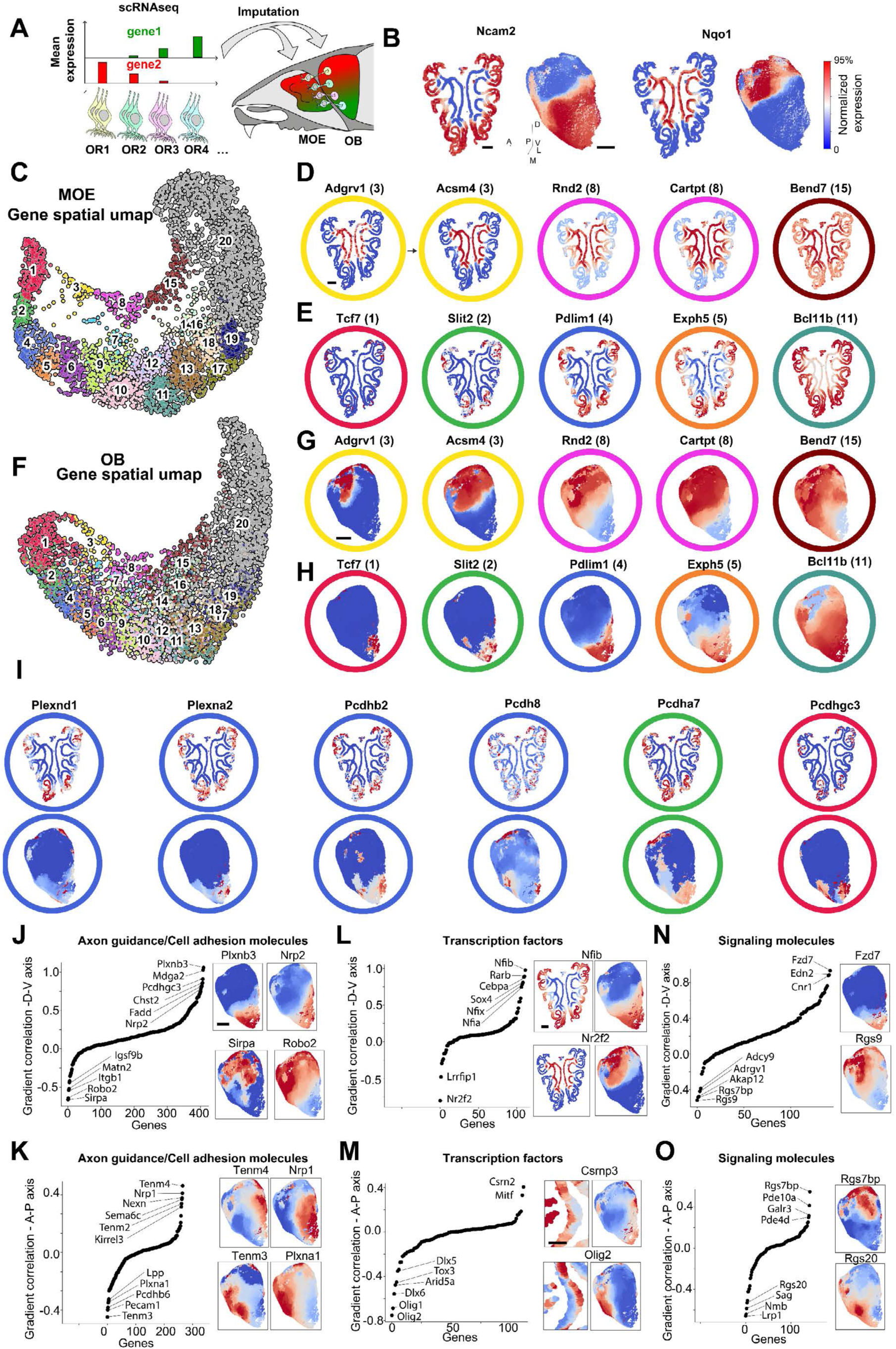
Integrating the spatial organization of OSNs with scRNAseq to reveal molecular patterns in the MOE and OB. **(A)** Schematic of the imputation method used to relate gene expression in OSNs measured by scRNAseq^47^ to their MOE spatial position and OB projection mapped by MERFISH. The average gene expression in scRNAseq data across each OSN type was transferred onto the OSNs within a reference MOE section and to their corresponding projections in the OB. Denoised distributions across the MOE and OB were obtained by averaging the transferred gene expression within a radius 50, 400 µm around each point on the surfaces of the two organs respectively. **(B)** Imputed expression patterns for two example genes (*Ncam2* and *Nqo1*). The expression patterns are normalized in the range 0(blue) to 1(red) with the maximum value capped at the 95th percentile of imputed expression across the surface. **(C)** UMAP representation of the imputed patterns in the MOE for the top ∼5000 differentially expressed genes across OSNs. Each gene pattern was embedded using distance metric based on correlation (i.e. genes which are close in the UMAP have a correlated spatial pattern in the MOE). Leiden clustering was used to group genes with correlated spatial patterns (marked with different colors and numbers). **(D)** Imputed MOE patterns of a few example genes across the top UMAP branch in (C) covering clusters 3,8 and 15. **(E)** Imputed MOE patterns of a few example genes across the bottom UMAP branch in (C) covering clusters 1, 2, 4, 5 and 11. **(F)** UMAP representation of the imputed OB patterns. Genes which are close in the UMAP space have a correlated spatial pattern across the OB. **(G),(H)** Images of the imputed OB patterns of the genes shown in (D) and (E) respectively. **(I)** Imputed patterns of example genes with similar MOE patterns but with more variable OB patterns. **(J)** Correlation between the spatial patterns of genes with a defined gradient along the dorsal-ventral (D-V) axis of the OB. Correlations for the axonal guidance/cell adhesion class of genes are shown. Genes with the strongest gradient along the D-V axis are marked. Right insets show the imputed patterns of example genes with strong gradients. **(K)** Same as (J), but along the anterior-posterior (A-P) axis. **(L),(M)** and **(N),(O)** Same as (J),(K) for transcription factor and signaling molecule gene classes, respectively. Scalebar indicates 500 µm except for (M) where it indicates 200 µm.

The top ∼5000 differentially expressed genes (excluding *ORs*) were imputed onto the MOE and OB spatial atlases. To systematically visualize groups of genes with similar spatial imputations, we performed dimensionality reduction using UMAP where genes with correlated spatial patterns were embedded in close proximity to each other (Figure 4C). This analysis revealed two continuous branches in the UMAP: the top branch included genes such as *Adgrv1*, *Acsm4*, and *Rnd2*, for which expression progressively expanded from the center to the periphery of the MOE (Figure 4D); the bottom branch included genes such as *Tcf7*, *Slit2*, and *Pdlim1,* which showed the opposite trend, with expression progressively expanding from the periphery towards the center (Figure 4E). Gene imputations across the OB formed a largely similar UMAP representation and clustering (Figure 4F-H) consistent with the close correlation observed between the spatial distribution of specific ORs in the MOE and their corresponding projection map in the OB (Figure 3).

In addition, a subset of genes had expression patterns that followed a different distribution scheme, forming distinct yet partially overlapping patches in the OB even when constrained to the same location on the MOE. For instance, multiple members of the plexin and protocadherin classes of axonal guidance molecules, with expression largely constrained to the same peripheral region of the MOE, formed multiple patches across different positions within the OB (Figure 4I). These results may reflect a subset of genes associated with the local refinement of OSN projections in the OB, as previously proposed for the protocadherin gene family^49^.

Gene ontology annotations^50^ were used to classify the differentially expressed genes, and three functional classes were further examined: axonal guidance/cell adhesion molecules, transcription factors, and signaling molecules. All three classes covered the entire UMAP spaces for both the MOE and OB (Figure S8A, B, C) and the expression profile of each class of genes was sufficient to predict the location of the OR projection with a median spatial resolution of ∼500 µm, with significantly more accuracy than upon randomly shuffling the OR identities (Figure S9A, B, C). This is consistent with the view that these classes of genes may be involved in the control of OR specification and OB projection by OSNs^51^.

For each class, the genes were sorted based on the strength of their expression gradient along the two main axes of OR projection into the OB: dorsal-ventral and anterior-posterior (Figure 4J-O). Among axonal guidance/cell adhesion molecules (Figure 4J, K), this analysis highlighted many of the previously established regulators of the murine OB projections such as *Nrp1*, *Robo2*, *Plxna1*, and members of the protocadherin family^49,52–54^ as well as genes which, while uncharacterized in *Mus musculus*, were involved in establishing the OB projections in other species including, for instance, members of the teneurin family from *Drosophila melanogaster* (Tenm2, Tenm3, and Tenm4)^55^. A similar analysis across transcription factors and signaling molecules (Figure 4 L-O) highlighted genes with the strongest gradients along the two axes. Among transcription factors we note *Nfia*, *Nfib* and *Nfix* shown to play a crucial role in establishing the zonal organization of ORs in the MOE^18^. The role of other transcription factors including members of the retinoic acid pathway (including Rarb) or members of the thyroid hormone pathway (including Nr2f2) remain to be further assessed. Among signaling molecules we identified members of the GPCR signaling pathway including regulators of G-signaling (Rgs7bp, Rgs9, Rgs20) and cAMP signaling (Pde10a, Pde4d), supporting the proposed role of *OR* G-coupled signaling in axonal projection into the OB^51^. Overall, this spatial analysis catalogues potential regulators of OSNs to OB projections or other spatially organized signaling pathways providing a foundation for future functional experiments.

### High-throughput measure of specific odor responses in the MOE

Single molecule FISH of the immediate early gene *Egr1* (Figure 5A) combined with MERFISH measurements enabled us to assess the response of OSNs to distinct olfactory cues. Briefly, the MOE of animals exposed to distinct olfactory cues were harvested and imaged to quantify OR and *Egr1* expression, hence revealing OSN identity and odorant-evoked neuronal activity as previously established for vomeronasal and olfactory chemosensory detection^36,37^.

**Figure 5.**
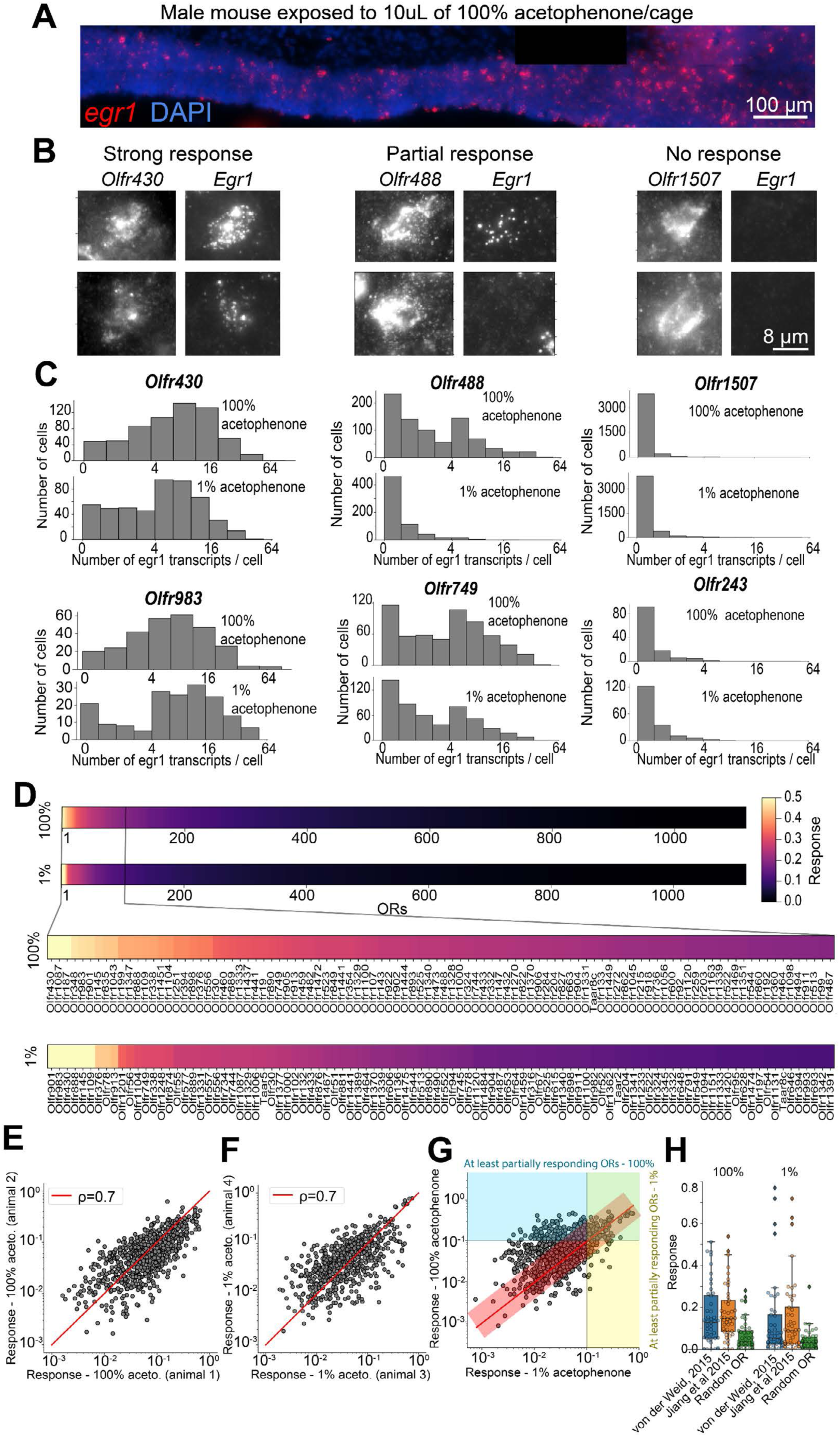
Cataloging OR responses to acetophenone. **(A)** Closeup of a small region of an MOE section from a male mouse exposed to 10uL of 100% acetophenone for 30 minutes. Single molecule FISH (smFISH) signal for *egr1* transcripts used a marker of neuronal activity^36^. **(B)** Two example sensory neurons expressing the same OR (*Olfr430* - left, *Olfr488* – center and *Olfr1507* – right) in which the OR and corresponding *egr1* signals are shown. *Olfr430* was selected as an example OR for which most OSNs of this type showed high levels of *egr1* expression upon 100% acetophenone exposure. *Olfr488* is an example OR for which only a fraction of the population of the corresponding OSNs express *egr1* (marked as partial response). *Olfr1507* is an example OR for which almost no OSNs are expressing *egr1* (marked as no response). **(C)** Histograms showing the distribution of the number of *egr1* transcripts per cell across example OSN types. The top and bottom histograms correspond to male mice exposed to 10uL of 100% acetophenone and 1% acetophenone, respectively. **(D)** Catalogs of OR responses to 100% (top) and 1% acetophenone (bottom). The response for each OR was quantified as the fraction of the population of neurons of a given OR-type which express >5 *egr1* transcripts. Responses were averaged across N=3 and N=2 male mice for 100% and 1% acetophenone respectively. **(E)** Correlation of OR responses between two male mice exposed to 10uL of 100% acetophenone. The Pearson’s correlation coefficient of 0.7 is marked in the inset. **(F)** Same as (E), for two male mice exposed to 10uL of 1% acetophenone. **(G)** Correlation between the OR responses to 100% vs 1% acetophenone. The red line marks equal response and the red transparent rectangle marks the region capturing 90% confidence interval based on the replicate-to-replicate correlation. Guidelines mark 10% response to 100% (horizontal line) and 1% acetophenone (vertical line). **(H)** Box and swarm plots of the responses of two sets of ∼50 ORs previously identified as responding to acetophenone (von de Weid et al 2015 and Jiang et al 2015). Left and right panels show responses to 100% and 1% acetophenone, respectively. Responses of a control group of 50 randomly selected ORs are displayed for comparison.

This paradigm was validated using the single-compound acetophenone, whose *in vivo* OR response was previously characterized by multiple methods^31,32^. Animals were exposed to two concentrations of acetophenone (100% and 1%, 10 µL each) applied to filter paper encapsulated in a permeable cassette placed in the cages of the animals. The number of *Egr1* transcripts detected in each OSN provided population-level responses for each OSN type (Figure 5B, C). OSNs expressing a given receptor displayed different types of responses: 1) OSNs expressing certain ORs (i.e. Olfr430 and Olfr983) had a *strong response* to acetophenone with the majority of the population co-expressing *Egr1* (Figure 5B, E, F), 2) other OSN types (i.e., expressing Ofr488 and Olfr749) had a *partial response* in which a subpopulation co-expressed *Egr1* (Figure 5G, H) and 3) most OSN types (i.e., expressing Olfr1507 and Olfr243) had predominantly no response with almost all corresponding OSNs lacking *Egr1* expression. The response of *ORs* to the cues was quantified based on the fraction of *Egr1^+^* OSNs (>5 *Egr1* transcripts/cell) for each OR type (Figure 5D and Table S6).

OR responses were quantitatively similar across different animals, for both acetophenone concentrations (Figure 5E, F, Person correlation coefficients of ∼0.7). *100%*- and *1%*-acetophenone concentrations also elicited correlated responses (Figure 5G, Pearson correlation coefficient of ∼0.6); however, a concentration-dependent shift was observed. Specifically, the number of strongly and partially responding ORs increased 2-fold upon increasing the concentration of acetophenone from 1% to 100%. Furthermore, 328 ORs had an increased response to *100%-* vs *1%*-acetophenone, beyond what is expected from the replicate-to-replicate variation in OR response (outside the 90% confidence interval). This demonstrates a concentration-dependent response at the population level of OSNs of different OR types.

Importantly, 7 of the 9 strongly-responsive ORs (∼80%) were previously identified by orthogonal sequencing approaches^31,32^ as responsive to acetophenone and, the other ORs identified in these studies showed statistically increased responses for both the high- and low-acetophenone concentrations compared to all other ORs (Wilcoxon test *p*-values of 8.4 × 10^−5^ and 9.8 × 10^−10^ for 100% acetophenone and Wilcoxon test p-values 3.6 × 10^−5^ and 4.4 × 10^−7^ for 1% acetophenone) (Figure 5H). These results suggest that our imaging approach is able to measure the response of the near complete repertoire of ORs to specific odor molecules.

### High-throughput measure of OSN responses to predator and social cues

MERFISH including *Egr1* imaging was performed on MOE harvested from animals exposed to different ethological cues, including predator (i.e. cat bedding) and social cues (male, female and pup intruders). OR responses were averaged across 2-3 animals for each exposure condition covering: 1) males exposed to cat odor as a predator cue, 2) females exposed to male intruders, 3) males exposed to male intruders, 4) males exposed to female intruders, 5) females exposed to female intruders, 6) virgin females exposed to pups and 7) mothers exposed to pups (Figure 6A and Table S7). As seen with acetophenone, graded responses were observed for both predator and social cues: a few ORs displayed a strong response (>50% of the OSNs expressing the same ORs co-expressed >5 *Egr1* transcripts per cell), 10-100 OSNs types displayed a partial response (defined as >10% of the OSNs expressing the same OR co-expressed > 5 *Egr1* transcripts per cell) and the rest of the OR repertoire showed no significant response (< 5 *Egr1* transcripts per cell) to the cue.

**Figure 6.**
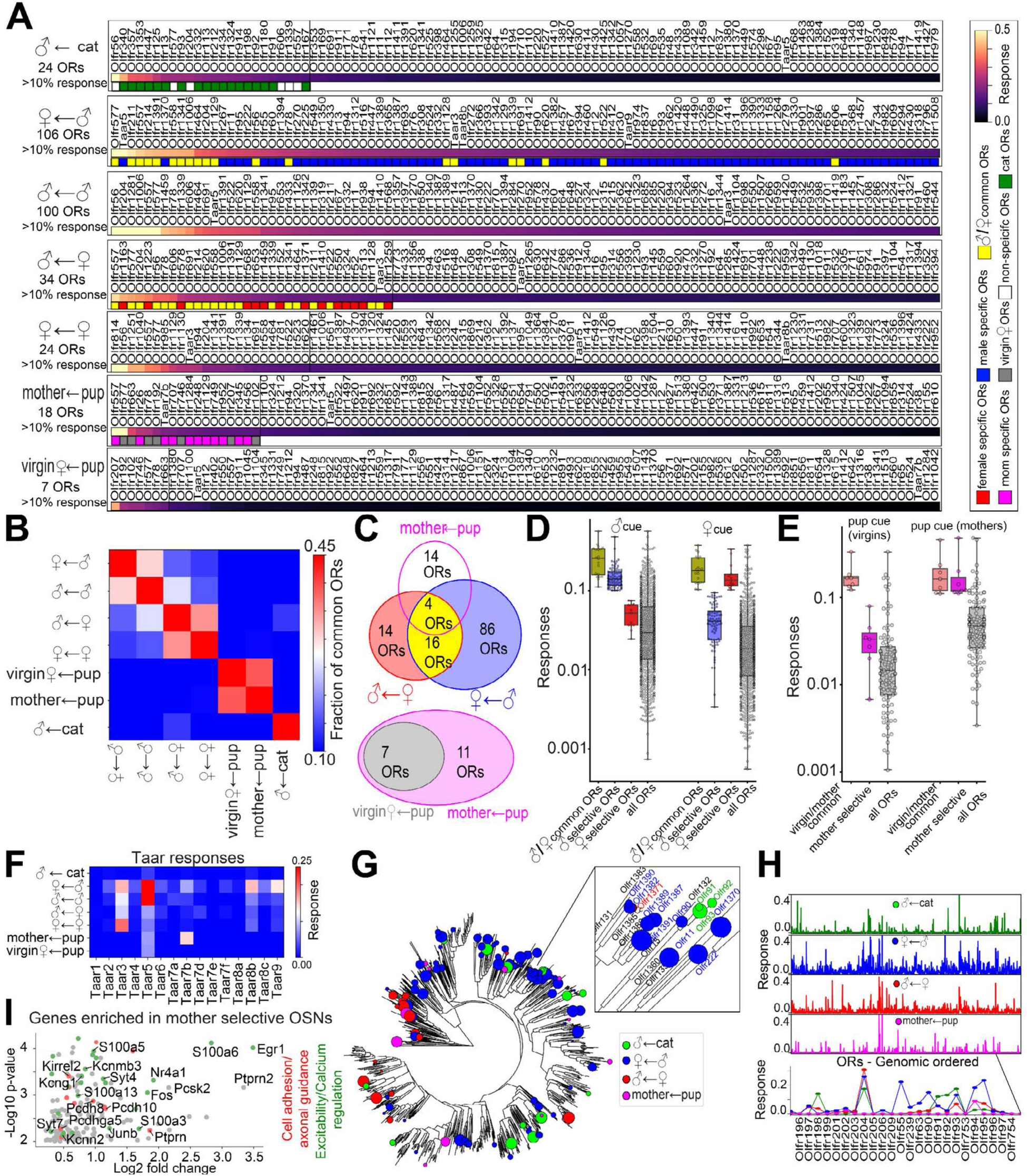
Cataloging OR responses to ethological cues. **(A)** Plot of responses of the top 100 ORs to the following olfactory stimulations: cat cue (sensed my males), male cue (sensed by females), male cue (sensed by males), female cue (sensed by males), female cue (sensed by females), pup cue (sensed by mothers) and pup cue (sensed by virgin females). For each plot the number of ORs with >10% response, defined as *responding* ORs, is indicated. The response of N=2-3 mice was averaged in each of these plots. Colored squares mark the following OR response types: white - non-selective ORs responding to the majority of cues, green - ORs selective to cat cue, blue - the ORs selective to male cue (sensed by females) compared to female cue (sensed by males), red - ORs selective to female cue (sensed by male animals) compared to male cue (sensed by female animals), yellow - ORs responding to both male and female cues (sensed by the opposite sex), magenta - ORs responding to pup cues selectively in the mothers compared to virgin females and gray - ORs responding to pups in virgin females. **(B)** Matrix showing the fraction of common responding ORs between each pair of the different exposure conditions labeled as in (A). **(C)** Top: Venn diagram of ORs responding to: male cue sensed by females (blue), female cue sensed by males (red), and pup odor sensed by mothers (magenta); Bottom: Venn diagram of ORs responding to pup odor sensed by the mothers (magenta) and by virgin females (gray). **(D)** Box and swarm plot of OR responses to male cue (sensed by females, left) and female cue (sensed by males, right) of different classes of ORs: 1) common male and female responding ORs (yellow), 2) male selective ORs (blue), 3) female selective ORs (red), and 4) all ORs (light gray). **(E)** Box and swarm plot of OR response to pup cue by virgin females and by mothers (right) of different classes of ORs: 1) common virgin and mother responding ORs (salmon), 2) mother selective ORs (magenta) and 3) all ORs (light gray). **(F)** Plot with the responses of Taars across the exposure conditions listed in (A). **(G)** OR phylogenetic tree annotated with responses to 4 olfactory cues: cat odor (sensed by males, green), male cue (sensed by females, blue), female cue (sensed by males, red) and pup cue (sensed by mothers, magenta). Dot size is proportional to OR response. Inset zooms-in on a clade containing subbranches Olfr1370-Olfr1390 and Olfr90-Olfr92 **(H)** OR responses to 4 olfactory cues with the ORs sorted along the genomic axis. Inset (bottom) zooms in on the responses of an example OR genomic cluster. **(I)** Differential gene expression plot comparing mother-selective OSNs to all other OSNs, based on scRNA-seq data^47^.

The responding sets of ORs, defined for each cue as having at least a partial response, were compared across the different cues and conditions. ORs showed high overlap in their response to the same cue (male, female, or pup) irrespective of the sex and parental experience (e.g. virgin females vs mothers) of the exposed animals (Figure 6B), indicating that these responses were primarily determined by the odorant itself rather than by the sex or internal state of the animal.

The number of responding *ORs* and their specificity varied across cues. ORs responding to male cues formed the largest and most selective set comprising 106 ORs out of which 19% also responded to female cue (Figure 6C). By contrast, ORs responding to female cues constituted a more reduced set (34 ORs), with 60% also responding to male odor to a similar degree. Notably, ORs categorized as male-selective (with >10% of corresponding OSNs co-expressing *Egr1* for male odor and <10% expressing *Egr1* for female odor) also had a statistically increased response to both sexes compared to all other ORs (p-value = 3.34 × 10^−5^ Wilcoxen test, Figure 6D). Nevertheless, their response was 3-4 times more highly tuned to male compared to female odor. Similar results were observed for the female selective ORs.

ORs responding to pup cues in both virgin females and mothers accounted for the most reduced set with only 18 ORs, of which 4 also responded to both female and male cues (Figure 6C). Strikingly, pup responses were modulated by the state of females: ORs responding to pups in mothers included all pup-responsive ORs identified in virgin females plus 11 additional, mother-selective ORs. ORs responding to pups in virgin females had approximately the same fraction of responding OSNs in mothers, while the mother-selective ORs had ∼4-fold higher response to pups in mothers compared to virgin females (Figure 6E). These data suggest that the increased response to pup cues in mothers originated from a subset of OSNs that become selectively more responsive to pups, while the response from other pup-sensitive OSNs does not change.

The response of Taars to ethological odors has been extensively investigated. Taar5 was identified as strongly responding to male cue in both male or female mice, with Taar5 deletion resulting in decreased sensitivity to male urine^56–58^. Consistent with these findings, our data indicate a strong response of Taar5 to male cues together with a modest response to female and pup cues (Figure 6F). For each of the other cues tested another member of the Taar family had the strongest response (Taar3 for female and Taar7b for pup cues) indicating that even across the narrower set of social cues and within the small family of Taars, receptor responses are differently tuned to specific ethological cues.

Although distributed across multiple clades of the phylogenetic tree, ORs responsive to a given ethological cue were largely more similar in sequence than expected by chance (Figure 6G, Wilcoxen test, *p*-values of 8.6 × 10^−8^, 3.0 × 10^−25^, 3.0 × 10^−190^ and 8.2 × 10^−3^ for cat, female, male and pup odors respectively). However, larger OR clades or genomic clusters showed a heterogeneous response (Figure 6G, H). For instance, the phylogenetic clusters Olfr1380-Olfr1390 and Olfr94-Olfr95 responded primarily to male and female cues while the neighboring cluster of Olfr90-Olfr92 responds primarily to cat cue. This heterogeneity is in contrast with the response of the vomeronasal V2R system^36^ in which the phylogenetic clades of different V2Rs were highly sensitive to specific predatory/social cues.

To investigate potential regulatory mechanisms underlying the increased response to pups observed in specific OSN types from mothers compared to virgin females, we searched for genes differentially co-expressed within these OSNs. OSN types showing selective pup responses in mothers were defined as OSNs with a response greater than 0.05 in mothers and with a more than two-fold higher response in mothers compared to virgin females. We searched for genes exhibiting significantly higher expression in these specific OSNs relative to all other OSN types in the sequencing dataset^47^. We restricted the analysis to cells coming from female animals, as classified on the expression of genes from X and Y chromosomes and identified 156 genes with significant differential expression (p-value < 0.01, Wilcoxon test), predominantly belonging to four functional categories: 1) axonal guidance molecules, including Kirrel2, Pcdh10, Pcdh8, Pcdhb18 and Pcdhga5, potentially reflecting the specialized projections of these OSNs; 2) regulators of neuronal excitability, including ion channel modulators (e.g., Kcng1, Kcnn3, and Kctd1), 3) regulators of calcium homeostasis including calcium-binding proteins (e.g., S100a3, S100a5, S100a6, and S100a13) and synaptotagmins (Syt4 and Syt7, and Syt9), and 4) immediate early genes (e.g., Fos and Junb) (Figure 6I).

Taken together, these experiments show a remarkable selectivity of OSN responses to ethologically relevant cues and activity modulation of specific OSNs in mothers.

### Integrating functional and spatial information onto the epithelium and olfactory bulb

The molecular identity of the OSNs defined by their OR expression allowed us to map olfactory responses onto the MOE and OB. OSNs expressing ORs responding to ethologically relevant olfactory cues were highlighted in a reference MOE slice to facilitate comparison across different cues (Figure 7A, B). Similarly, the projections of the responding OSN sets were highlighted in a 3D OB projection map (Figure 7C, D). The different classes of cues profiled corresponded to distinct spatial distributions both within the MOE and the OB. For example, receptors responding to cat cue were primarily enriched in a more peripheral circle within the MOE (Figure 7A) and projected primarily to a thin ventral band within the OB with a higher density at the most anterior and most posterior regions (Figure 7C, E, F). ORs responsive to male odor were more broadly distributed across the MOE and projected primarily to two regions of the OB - a small dorsal region and a large mid-band along the dorsal-ventral axis, spanning most of the anterior-posterior direction. ORs responding to female odor also covered a large area of the MOE, with preference to the central region, and projected to a few OB subdomains along the dorsal-ventral axis and more restricted to the middle of the anterior-posterior axis. Finally, ORs responding to pup cues were concentrated primarily in the central MOE and projected to a small dorsal subdomain of the OB with a few scattered projections corresponding to ORs with a lower pup response. Notably, the dorsal subdomain covered the most responsive ORs in mothers compared to virgin females. These data begin to define a molecular and spatial organization of responsive OR domains within the MOE and OB and reveal the spatial specificity of ethologically relevant odor responses.

**Figure 7.**
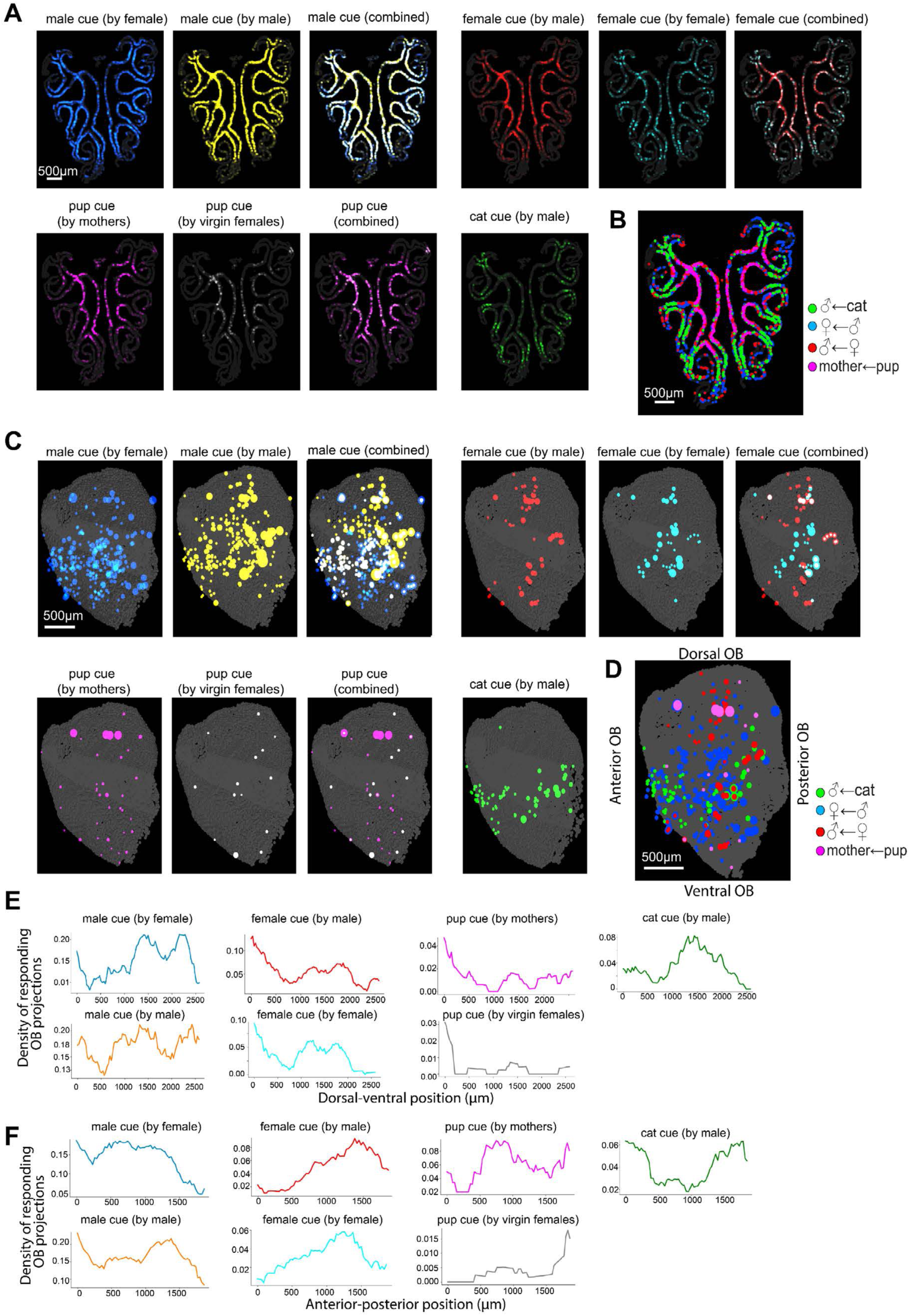
The spatial distribution and projection of the ORs responding to different olfactory cues. **(A)** Images of a reference coronal section of the MOE with OR responses mapped to their corresponding OSNs. A transparency scale was used for each depicted OSN proportional to its corresponding OR’s response. **(B)** Composite image summarizing MOE spatial patterns responding to different cues. **(C)** Images of olfactory bulb projections mapped based on their OR response to different olfactory cues. Dot sizes are proportional to OR responses. **(D)** Composite image summarizing the different OB spatial patterns of responses to different cues. **(E)**,**(F)** Distribution of responding OR projections for each of the cues along the dorsal-ventral (D-V) and anterior-posterior (A-P) axis of the OB, respectively. The density of responding OR projections was defined along each axis by binned in 200µm intervals.

#### Discussion

This study adopts a multiplexed imaging approach to systematically determine expression across the near entire OR gene family both within the somas of sensory neurons and their axonal terminals. Two high resolution and comprehensive molecular atlases were constructed capturing the spatial distribution of sensory neurons within the MOE and their 3D projections into the OB. In the MOE, the dominant mode of organization of ORs was the formation of continuous rings of increasing size from the center to the periphery. ORs displayed an additional smaller scale organization along the apical-basal direction perpendicular to the MOE turbinates. In the OB, the OR projection map was stereotypical both between bilateral bulbs and between bulbs of different animals with an average precision of axonal targeting of 200-300 µm. This result aligns with prior work^19^ showing that individual glomeruli can be matched across animals by their odor-response fingerprints with a positional variability of ∼200– 300 µm (the size of 1-2 glomeruli), underscoring the high spatial and molecular precision of the olfactory bulb map. An intriguing geometrical connection emerged upon comparing the MOE and OB atlases: the preferential distribution of ORs along the central-peripheral and the basal-apical axes in the MOE was remapped onto the dorsal-ventral and anterior-posterior axes in the OB, respectively.

These atlases represent a valuable resource and provide insights into genes that orchestrate the intricate and stereotypical spatial organization of sensory neurons and their projections into the OB, when integrated with existing or future molecular and functional studies. For instance, in this study, taking advantage of the availability of the ever-increasing quality of the single-cell RNA sequencing datasets of sensory neurons^47,59,60^ through imputation methods, we began a preliminary investigation of the interplay between differential gene expression across OSNs, their precise spatial location in the MOE and their axonal projection in the OB. Many genes across different families of transcription factors, axonal guidance and signaling molecules appeared to be expressed within neurons projecting to specific, contiguous subregions of the OB glomeruli layer. Furthermore, the expression profile of each of these gene classes in the OR neuronal types had a high predictive power for the 3D location of the projection of the corresponding OR type in the OB (Figure S8). These results support the notion that transcription factors and signaling molecules work together to synchronize the OR expression with the expression of appropriate signals for axonal guidance.

Placing the spatial organization of sensory neurons within their functional context is a particularly exciting direction, which will synergize and further strengthen newly developed methods to de-orphan receptors^31,32^. Many of these methods test monomolecular odorants at high concentration but often lack diversity represented in ethologically relevant odor mixtures. Hence, we leveraged the sensitivity and throughput of the MERFISH technology coupled with smFISH for immediate early gene expression (*Egr1*) to functionally characterize the OR responses to different social and predator cues. The specific sets of responding ORs to each cue were catalogued and spatially mapped within the molecular atlases of the MOE and OB. This revealed the distinct spatial organization and projection of the responsive ORs within the MOE and OB, suggesting that odor information is mapped within constrained regions for localized processing. The spatially localized response to male odor during direct interaction aligns with previous studies mapping IEG expression in the OB upon male urine exposure^61^. Elevated activity was found in OSNs projecting to a central band along the dorsal ventral axis of the OB and, to a lesser extent, to a dorsal subregion. By contrast, responses to female odor were more enriched in OSNs projecting to the dorsal OB, which has been previously shown to be functionally critical for generating responses to female cues compared to male cues^62^. Prior efforts mapping IEG expression upon exposure to cat predatory odor reported only a modest increased response in the OB, limiting precise spatial characterization of the odor response^63^. Here, we found that the response to cat predatory odor was localized primarily to OSNs projecting to a thin ventral band encircling the OB. This finding, combined with the enrichment of pup sensitive ONSs in the dorsal region of the OB, challenges a prior hypothesis that ventral regions of the OB encode attractive cues while the dorsal regions correspond to aversive cues^1^. The response to pup cue further revealed an increased sensitivity in a subset of OSN types in mothers compared to virgin females. While the prolactin signaling, whose modulatory role in females was recently described^64^, was not found to be enriched in mother selective OSNs (not shown), we found other modulatory genes enriched in these OSNs involved in ion channel and calcium regulation. We note, however, that the number of cells in the single-cell sequencing data used is modest (∼10,000 female OSNs) and hence certain modulatory genes might have been missed from this analysis. We also note that apart from transcriptional differences across OSNs, other factors could contribute to modulating OSN responses including, for instance, the animal’s sampling duration when probing different ethologically relevant odors, the kinetics of the ORs and their downstream signaling cascade. The identification of OSNs that selectively detect social and predator cues paves a new avenue to study the processing of distinct categories of innate main olfactory cues associated with dramatically different behavioral repertoire.

As the OR responses of more cues get characterized using either the imaging approach introduced here, or other molecular/imaging approaches, we anticipate that the spatial atlases provided here will help understand how the chemical environment gets progressively mapped across the olfactory circuit including the epithelium and the olfactory bulb.

#### Limitation of the study

Mechanisms orchestrating the intricate spatial organization and projection of olfactory sensory neurons are still unclear. Nevertheless, we anticipate that the comprehensive atlases of OSN distribution and projection presented here will help address this question when integrated with future molecular and functional studies. For instance, our preliminary exploration of the imputations of gene expression from single-cell sequencing onto the OR spatial atlases highlighted many axonal guidance molecules whose roles in OSN axonal guidance have been established. However, identifying transcription factors and signaling molecules involved in OSN projections or constraining OSNs within specific MOE locations is an emerging, yet incomplete effort^18^ which will be aided by the data provided in this work.

Connecting the spatial organization of sensory neurons with their role in sensing different ethological odors is incomplete. Here we catalogued responses to one predatory cue (cat) and three social odors (male, female and pups). Future studies applying and extending the methodologies introduced here will continue cataloguing other classes of cues including different classes of food odors, of nesting materials or of different predators and provide a more comprehensive understanding of how odor information is mapped across the olfactory circuitry.

We further note that IEG expression used in this study is only an indirect proxy for neuronal activity. An alternative, more direct approach to measure the neuronal activity of sensory neurons in response to different olfactory cues is to fluorescently monitor calcium changes over time. Two-photon microscopy in transgenic lines expressing activity reporters^36^ enabled the live recording of the Ca^2+^ activity over time in the dorsal layer of glomeruli as head fixed animals are presented to different cues. Considering the recent advances in recording of Ca^2+^ activity within sensory neurons within the chemosensory organs^65^, we envision that after the live recording, the animals’ OBs or MOEs can be dissected for MERFISH imaging of OR genes. While these experiments are technically challenging and can only provide access to a limited volume of the MOE/OB, they would provide a more direct connection between the functional and molecular data complementing the more comprehensive IEG-based measurements.

## ACKNOWLEDGEMENTS

This work was in part supported by the National Institute of Mental Health (U19MH114821 to X.Z. and C.D., R01HD082131 to C.D., DP5-OD030878 to B.B.). X.Z. and C.D. are HHMI investigators. We thank S. Sullivan for help with animal exposure to different cues, MCB Graphics from Harvard University for helping with figures, Drs. Murphy, Kaplan and Talay for helpful comments on the manuscript. We also thank Dr. L. Tan for helpful discussions in designing the MERFISH probes targeting olfactory receptors.

## DECLARATION OF INTEREST

X.Z. is an inventor of patents applied for by Harvard University related to MERFISH. X.Z. is a co-founder and consultant of Vizgen, Inc. The other authors declare no conflict of interest.

## Figures

**Figure S1.**
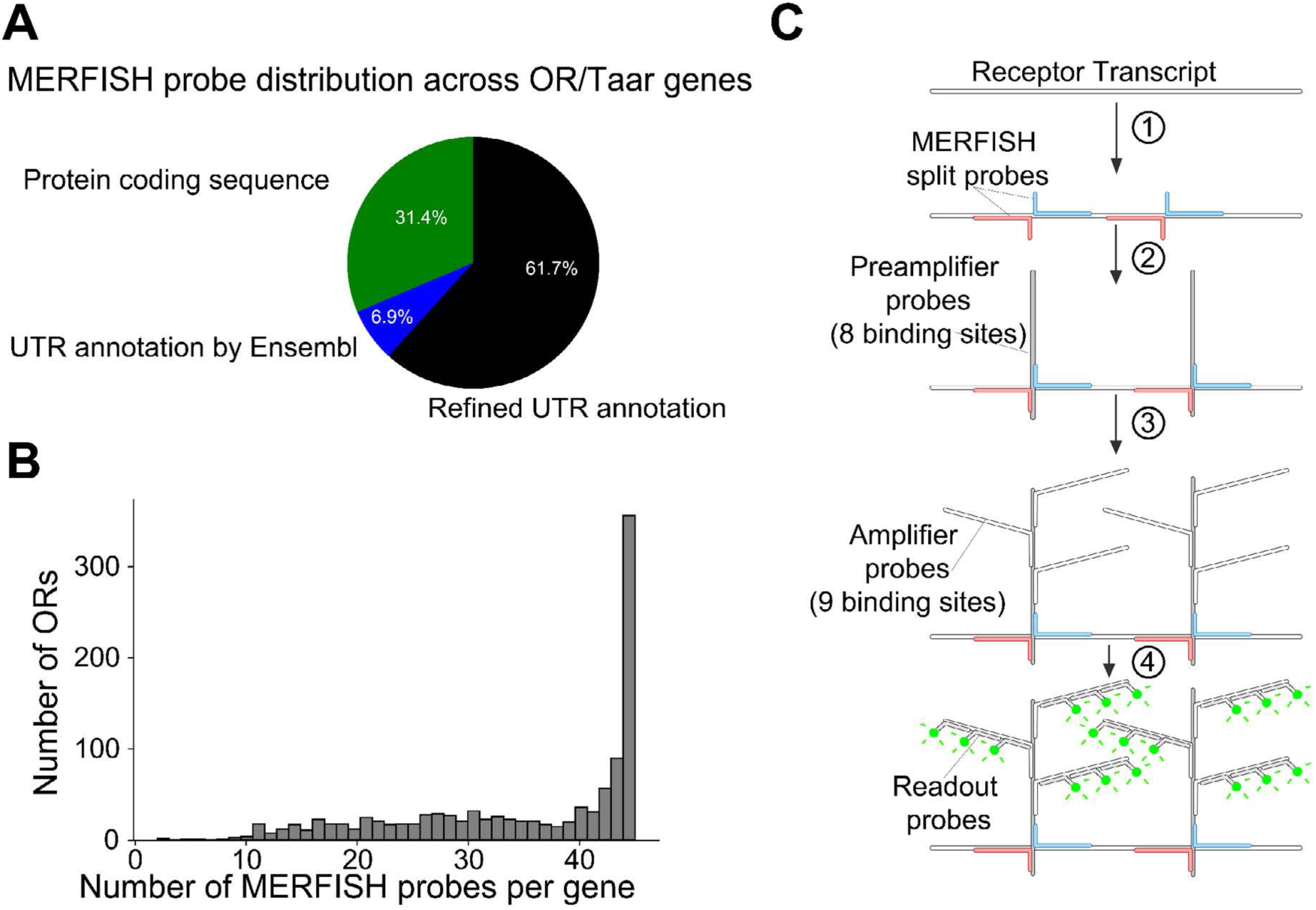
Modification to the MERFISH probe design to enable genome-scale OR/Taar targeting. **(A)** Pie chart with the percentage of the MERFISH encoding probes targeting the receptor coding region (green), the Ensemble annotated UTR regions(blue) and the extended annotation (from ^40^) of the receptor UTR (black). **(B)** Histogram with the distribution of the number of split-pairs of encoding probes across OR/Taar genes. **(C)** Schematic of the split-branched amplification scheme. In the 1st step pairs of encoding oligonucleotide probes are first hybridized to the receptor transcripts. In the 2nd step each pair of encoding probes colocalized onto the transcript allows the stable binding of a 200-nt single stranded DNA probe, called a preamplifier probe. The preamplifier contains eight 20-nt repetitive binding sequences. In the 3rd step, a similarly designed 200-nt single stranded DNA probe, called an amplifier probe, binds to the repetitive binding sites of the preamplifier. Finally in the 4th step, 30-nt fluorescent oligos called readout probes bind the repetitive sequences of the amplifier probes.

**Figure S2.**
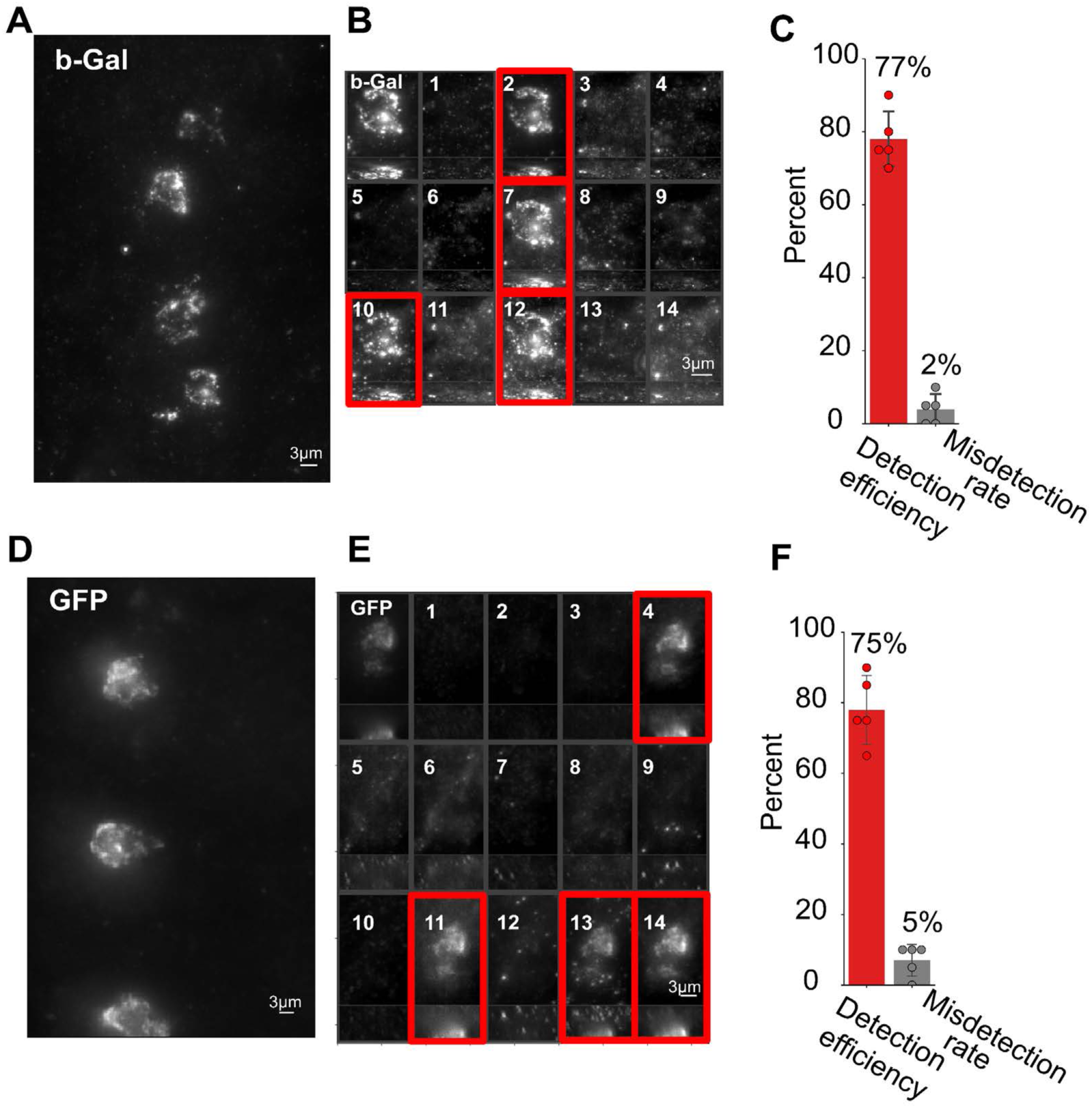
Quantifying the detection efficiency and accuracy of MERFISH using transgenic lines. **(A)** An image with the fluorescent signal of smFISH probes hybridized to β-Gal in a section of the MOE of a Olfr17-IRES-tau-lacZ transgenic animal. **(B)** Max-projection images of a β-Gal positive neuron showing the signal of the β-Gal probes (top left panel) and the MERFISH signal across 14 readout cycles. Readouts 2, 7, 10 and 12 (highlighted in red) correspond to the 4 readout probes binding to the MERFISH encoding probes designed for the Olfr17 transcript. **(C)** Bar plots marking the detection efficiency and misdetection rate for the Olfr17 transcript. The detection efficiency is defined as the fraction cells expressing β-Gal identified as Olfr17 positive via MERFISH. The misdetection rate is defined as the fraction of cells identified as Olfr17 positive via MERFISH which do not express β-gal. Quantification was performed across 2 MOE sections. The mean and standard deviation are plotted. **(D),(E)** and **(F)** Same as (A), (B) and (C), respectively, for Olfr16 in an Olfr16-IRES-tau-GFP transgenic animal.

**Figure S3.**
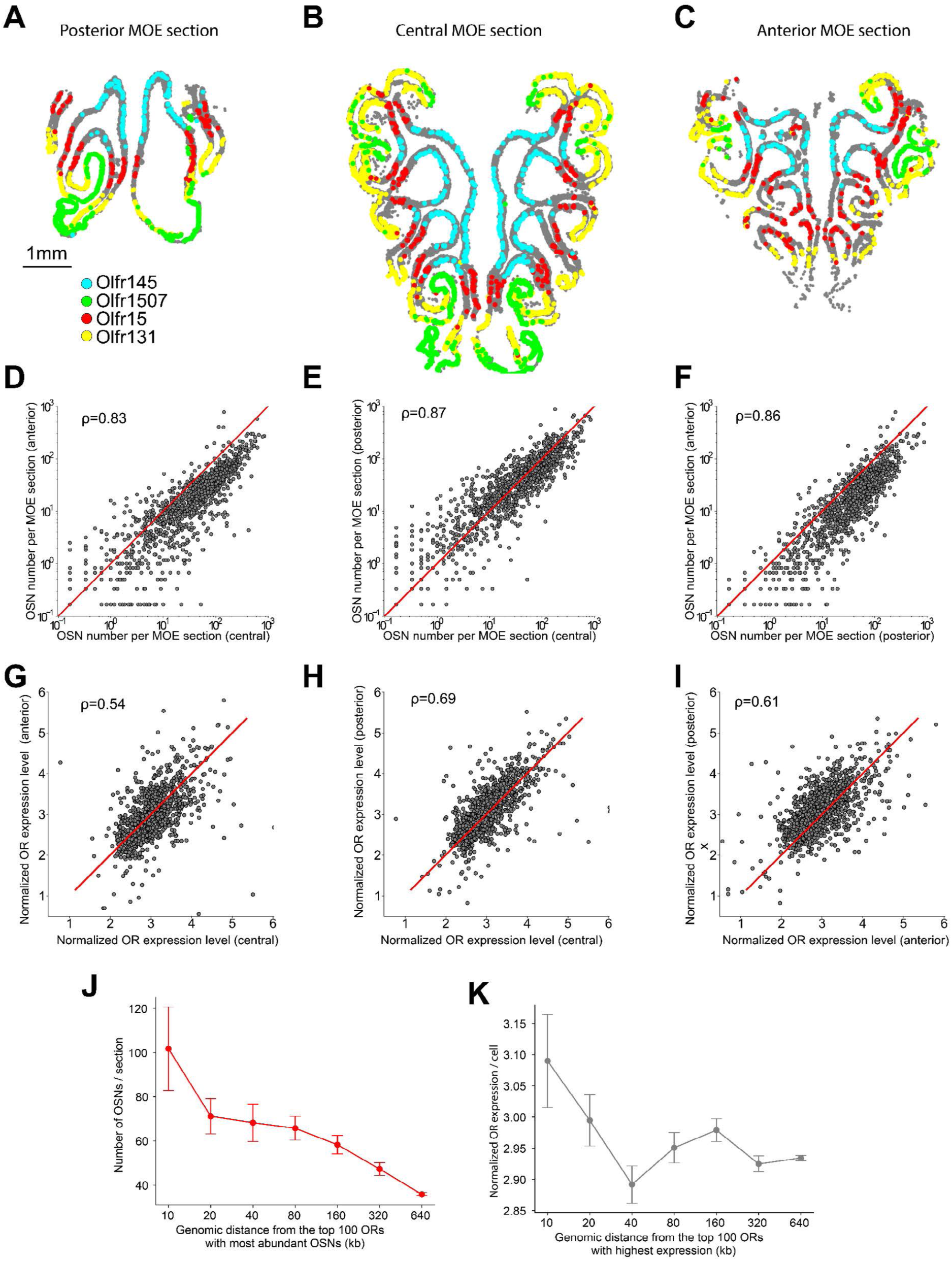
OSN abundance and OR expression level per cell are similar across the anterior-posterior axis of the MOE. **(A), (B), (C)** Representative coronal sections from the posterior, central and anterior MOE, respectively, overlayed with OSN positions of 4 example ORs. **(D), (E), (F)** Correlations between the number of OSNs per section for each OR comparing anterior to central, posterior to central and anterior to posterior MOE sections respectively. **(G),(H), (I)** Same as (D), (E), (F) for the average OR expression for each OSN type. **(J)** Correlation between the average number of OSNs per section and the genomic distance from the top 100 ORs with most abundant OSNs. **(K)** Correlation between the average OR expression per cell and the genomic distance from the top 100 ORs with highest expression. Error bars mark the standard error of the mean.

**Figure S4.**
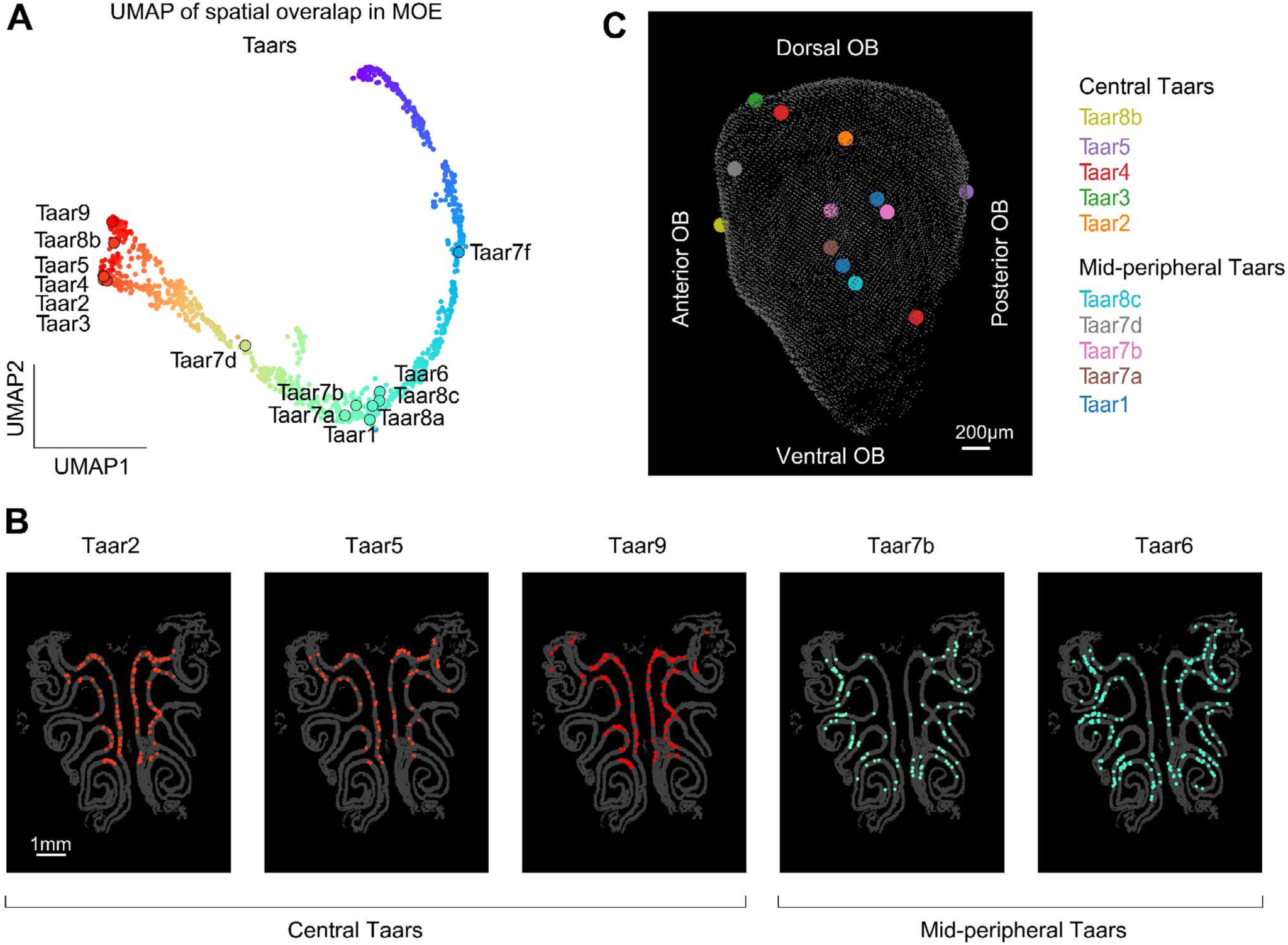
The spatial distribution of Taars within the MOE and OB. **(A)** UMAP representation of ORs based on their spatial overlap (reproduced from Figure 2B) which Taar genes overlayed. Taars are primarily located central or mid-peripheral MOE **(B)** Example images showing the spatial distribution of 3 Taar genes within the central MOE and 2 Taar genes within the mid-peripheral MOE. **C.** Image highlighting the Taar projections detected within the OB.

**Figure S5.**
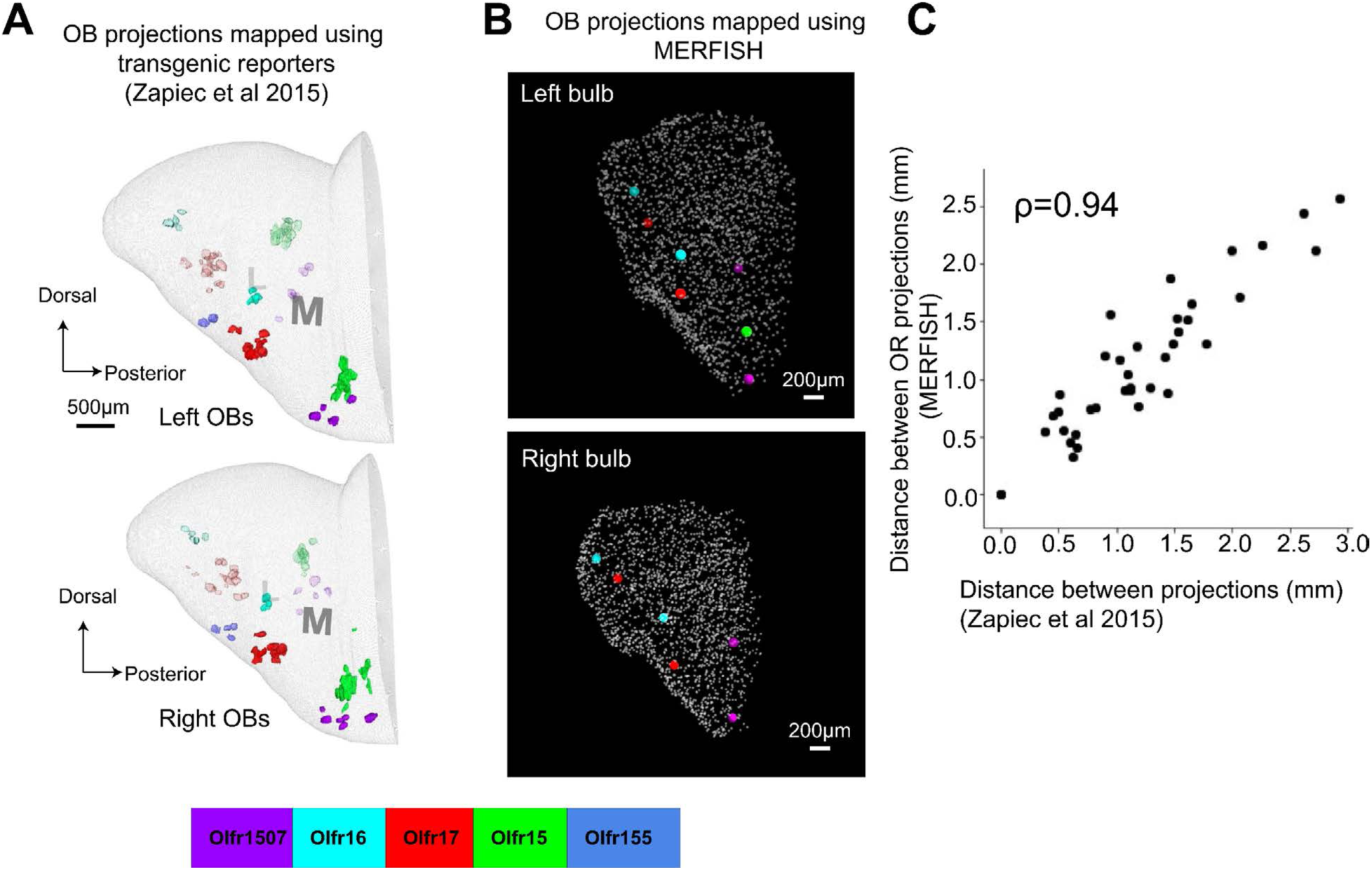
Comparing MERFISH identified OR projections with OR projections labeled in transgenic lines. **(A)** The 3D projections for 5 ORs identified by serial two-photon tomography in transgenic lines with fluorescently labeled ORs. The left and right bulbs of multiple animals were aligned to construct a unified map. This panel is reproduced from^20^. **(B)** MERFISH identified projections for the corresponding ORs across the left and right OB of a CD1 female mouse. **(C)** Correlation between the pairwise distances of OR projections identified in ^20^ and the corresponding pairwise distances of OR projections identified by MERFISH. Pearson correlation coefficient of 0.94.

**Figure S6.**
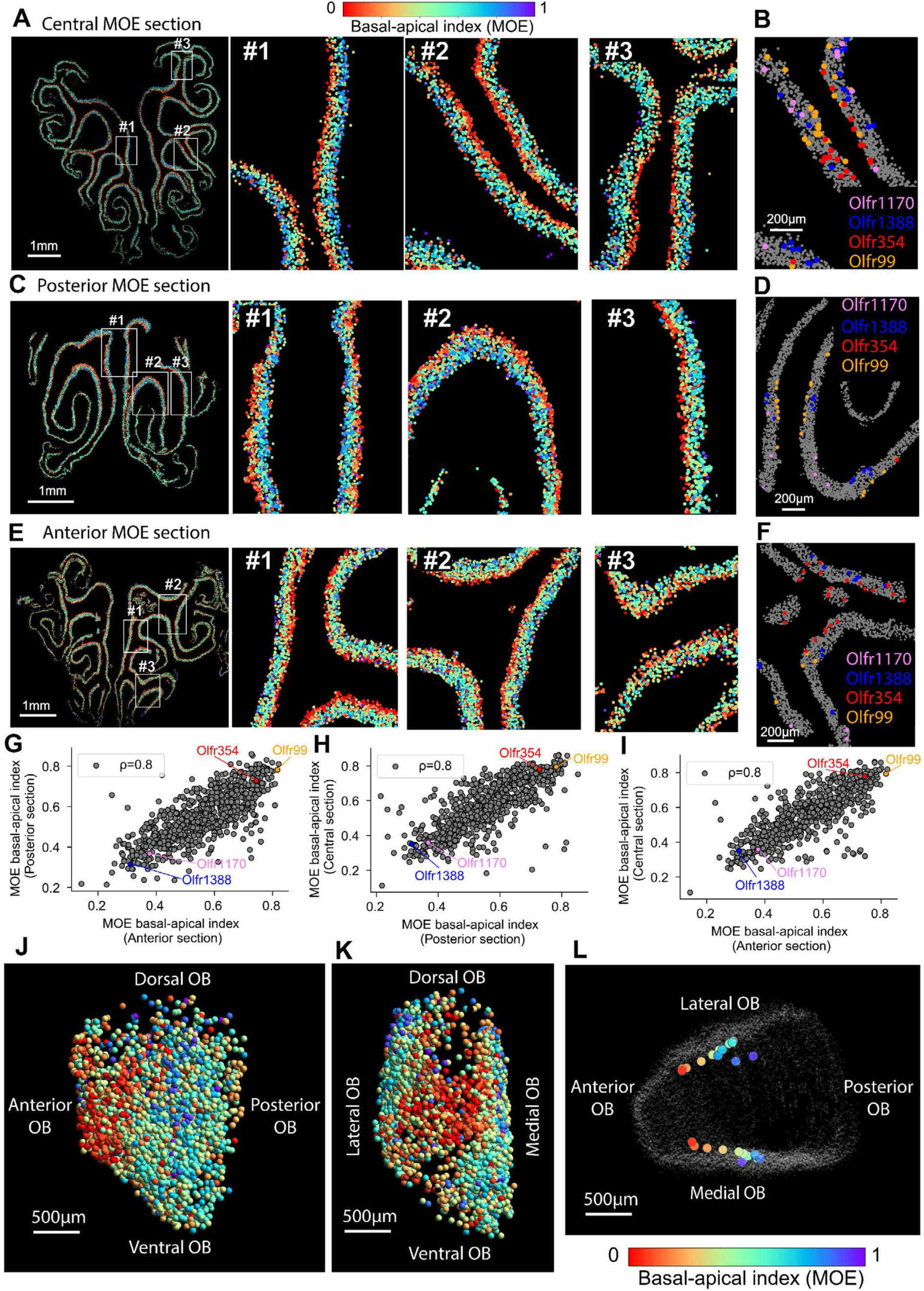
The basal-apical position of OSNs in the MOE correlates with the anterior-posterior position of their projections in the OB. **(A)** Example image of a central MOE section with OSNs labelled based on the average basal-apical position of each OR type. Zoom-ins are shown for 3 different MOE regions. **(B)** Zoom-in image on a MOE region with the positions of 4 example OSN types highlighted (Olfr354 and Olfr99 are more basal while Olfr1388 and Olfr1170 are more apical). **(C),(D)** and **(E),(F)** same as (A),(B) for a posterior section and an anterior MOE section respectively. **(G),(H),(I)** Pairwise correlation of the basal-apical position of OSNs between anterior, posterior and central sections of the MOE. **(J)** OB projections colored based on the basal-apical position of their corresponding OSNs in the MOE (medial side). **(K)** Same as (J) shown from a different view angle of the OB. **(L)** Image of the OB together with the average positions of OR projections binned based on the basal-apical positions of the corresponding OSNs in the MOE. Two parallel axes (one for medial projections and one for lateral projections) emerged along the anterior-posterior axis of the OB.

**Figure S7.**
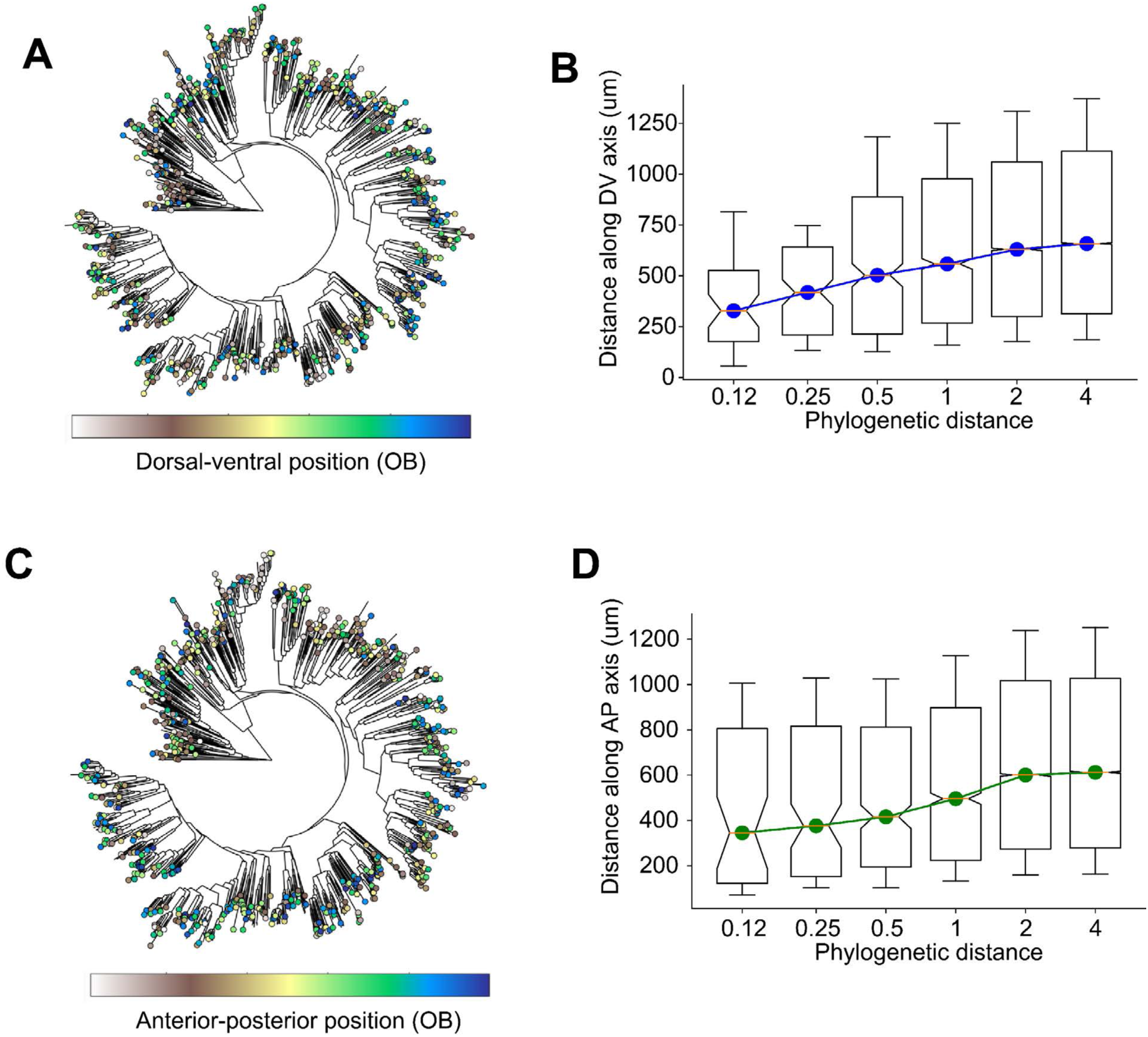
Relationship between OR phylogenetic distances and OB projections along dorsal-ventral and anterior posterior axes. **(A)** Phylogenetic tree connecting *ORs* based on their sequence similarity, constructed as in^32^. Each node, representing an OR, is colored based on the dorsal-ventral position of the OB projection as indicated in the colormap. **(B)** Correlation between distances of *ORs* along the phylogenetic tree of pairs of ORs and distances along the dorsal-ventral projection axis. For each pair of ORs we calculated their distance along the tree in (A) and then binned these distances. For each bin we represent a box plot marking the median dorsal-ventral distances between the ORs within the corresponding phylogenetic distance range. Notches represent 95% confidence intervals, the boxes mark the first and third quartile and the whiskers mark 15th and 85th percentiles. **(C), (D)** Same as (A) and (B) respectively, for the anterior-posterior axis of the OB.

**Figure S8.**
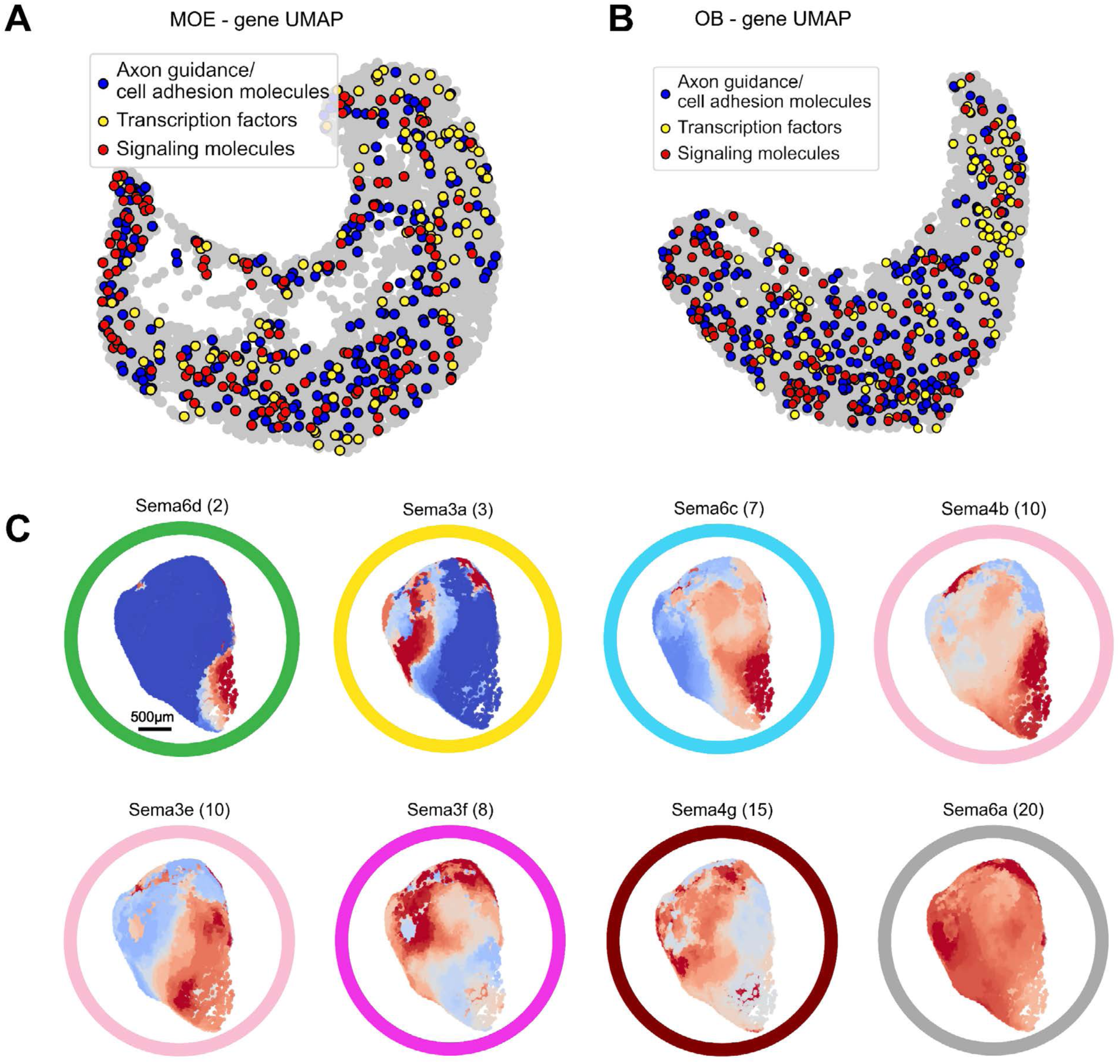
Variability of expression patterns imputed within MOE or OB for specific classes of genes. **(A)** UMAP representation of the MOE imputed spatial patterns (reproduced from Figure 4D) with the axonal guidance/cell adhesion molecules, transcription factors and signaling molecules overlayed. **(B)** Same as (A) for the OB UMAP reproduced from Figure 4F. **(C)** The imputed spatial patterns of the Semaphorin gene family in the OB.

**Figure S9.**
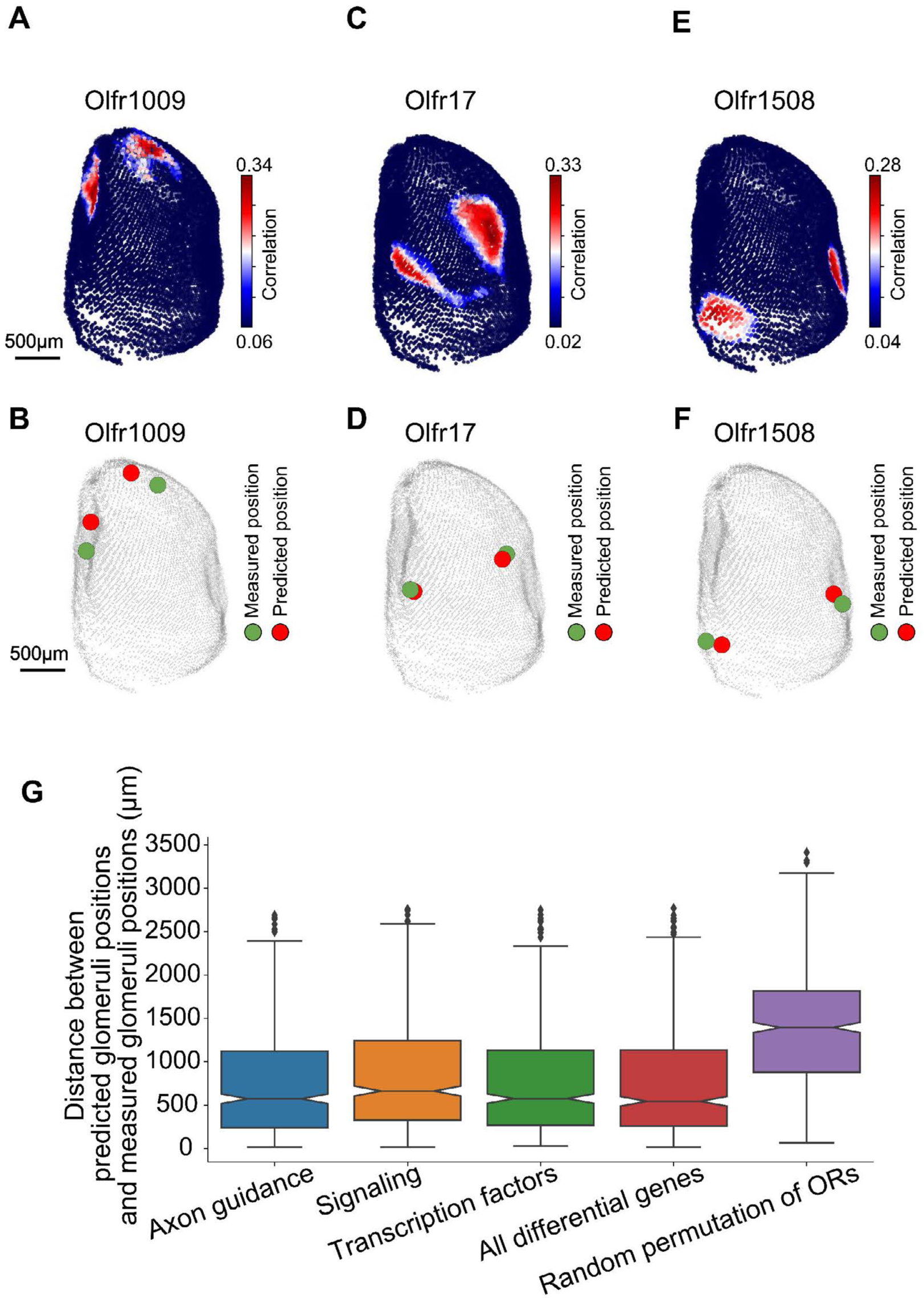
The predictability of OR projections in the OB based on the gene expression of OSNs in the MOE. **(A)** Correlation between the average gene expression of Olfr1009 expressing OSNs and the imputed expression across the OB from all the other ORs (excluding Olfr1009). The correlation was computed across the top ∼5000 differentially expressed genes for all points sampled on the glomerular layer. **(B)** OB image showing the projections of Olfr1009 measured by MERFISH (green) and the two points with the maximum medial and lateral correlations computed in (A) (red). **(C),(D)** Same as (A) and (B) for Olfr17. **(E), (F)** Same as (A) and (B) for Olfr1508. **(G)** Box plots showing the distances between the predicted glomeruli positions (red dots in (B), (D), (E)) and the MERFISH-identified glomeruli positions across the OR repertoire (green dots in (B), (D), (E)) when using different sets of genes for computing the correlation described in (A). Purple box plot shows the distances between the predicted glomeruli positions and the measured glomeruli positions using the correlation of the 5000 differentially expressed genes upon randomly shuffling the OR identities of the projections.

## Supplemental Table Legends

**Table S1. Sequences of the MERFISH probes and associated primers/readouts used to target olfactory receptor, trace amine-associated receptor and immediate early genes.**

**Table S2. The median number of OSNs per section and the median normalized OR expression per OSN across female and male animals.**

**Table S3. The central-to-peripheral index in the MOE of the OSNs detected and their coordinate on the spatial overlap UMAP.** The positions of all OSNs are also provided for 3 representative MOE sections from 3 animals.

**Table S4. The 3D positions of all the glomeruli whose receptor identity was determined by MERFISH.** Four olfactory bulbs were aligned in a single reference and the positions across them were concatenated. 3D positions defining the glomeruli layer surface of the reference olfactory bulb is also provided.

**Table S5. The basal-to-apical index of each OSN type.** Three MOE sections across different positions on the anterior-posterior axis were used to quantify the average basa-to-apical position of each OSN type. These positions were normalized such that 0 is the most basal position identified and 1 is the most apical.

**Table S6. The response of each OSN type (defined as the fraction of OSNs of each type containing >5EGR1 transcripts per cell) for animals exposed to two concentrations of acetophenone.**

**Table S7. The response of each OSN type for animals exposed to different ethological cues.**

## STAR Methods

### RESOURCE AVAILABILITY

#### Lead contact

Further information and requests for resources and reagents should be directed to lead contact, Catherine Dulac and first author, Bogdan Bintu.

#### Materials availability

Oligonucleotide probe sequences used for MERFISH imaging can be found in Tables S1. Templates and reagents for making these probes can be purchased from commercial sources, as detailed in the Key Resources Table.

#### Data and code availability

● Data reported in this paper are available at the Brain Knowledge Platform and Brain Image Library (https://knowledge.brain-map.org/data/KD2K0QPHBRWTNKYDUKB and https://knowledge.brain-map.org/data/7GQI8P3P8R40B6LPV9R/summary).
● Analysis code used for MERFISH probe library design, MERFISH image decoding, and data quantification is available at: https://github.com/ZhuangLab/MERlin, on Zenodo^66^ and https://github.com/BogdanBintu/MERFISH_Analysis_Olfactory_Receptors.
● Any additional information required to reanalyze the data reported in this paper is available from the lead contact upon request.

### EXPERIMENTAL MODEL AND SUBJECT DETAILS

#### Animals

C57BL/6J and CD1 mice were obtained from Jackson Laboratories and Charles River Laboratories respectively. Olfr17-IRES-tau-lacZ and Olfr16-IRES-tau-GFP mice were obtained from Dr. Richard Axel’s lab. The age, sex and sexual experience of each animal used is detailed below. Mouse husbandry and experiments were performed following institutional and federal guidelines and were approved by Harvard University’s Institutional Animal Care and Use Committee.

### METHOD DETAILS

#### Oligonucleotide Probe design

The design of the MERFISH probes targeting the olfactory receptor genes followed a similar design as previously described^67^ with a few notable modifications.

Briefly, the probes were synthesized from a pool of oligonucleotides purchased from Twist Biosciences. Each oligo in this pool consisted of the following sub-sequences (from 5’ to 3’):

1. A 20-nt or 19-nt forward priming region for PCR amplification
2. A 20-nt priming sequence used for the RT reaction during synthesis
3. A 15-nt 5’ split readout sequence used for developing the MERFISH signal
4. A 30-32-nt target sequence complementary to the olfactory receptor transcripts
5. A 15-nt 3’ split readout sequence used for developing the MERFISH signal
6. A 20-nt or 19-nt reverse priming sequence for PCR amplification

The target sequences were also screened using a similar procedure as in^67^ with the additional feature that we allowed the length to vary between 30-32 nt to increase the number of neighboring pairs of probes. Moreover, to increase the specificity of the target sequences to the OR of interest, we designed probes against the enlarged annotations of the olfactory receptor transcripts^40^.

#### Assigning the MERFISH readout sequences using a split-amplifier strategy

For every pair of adjacent probes targeting the OR transcripts we assigned a unique MERFISH bit by first designing a specific 5’ split readout sequence to the 3’ probe in the pair and a 3’ split readout sequence to the 5’ probe in the pair (see Figure S1 for details). The 3’ and 5’ pair split readout sequences together formed a single 30-nt readout binding sequence when the pair of probes are in proximity (i.e bound to the target transcript). A total of 15 pairs of 3’-5’ split readout sequences were designed - each corresponding to 1 of the 15 MERFISH bits. A unique set of 4 out of 15 3’-5’ pairs of split readout sequences were associated with the probes targeting each OR transcript. Each of these sequence pairs were repeated for each transcript 4-10 times.

190-nt probes (IDT), called preamplifier probes, were designed to contain a 30-nt sequence complementary to the concatenated pair of split readout probes. The remaining sequence contained 8 repeats of a 20-nt binding site. A total of 15 preamplifier probes were uniquely designed and correspond to each of the 15 MERFISH bits.

200-nt probes (IDT) called amplifier probes were designed to contain a 20-nt sequence complementary to the repeated binding site on each preamplifier probe and 9 repeats of another 20-nt binding site. A total of 15 preamplifier probes were uniquely designed to bind to each of the preamplifier probes and correspond to each of the 15 MERFISH bits.

30-nt fluorescent readout probes (IDT) were designed to contain a 20-nt sequence complementary to the repeated binding site on the amplifier probe. 15 readout sequences were designed to bind each amplifier probes and corresponded to each of the 15 MERFISH bits.

The *Egr1* probes were designed in a similar manner with a unique pair of 3’-5’ split readout probes and unique preamplifier, amplifier and readout probes. The Egr1 signal was first labelled and imaged across the sections, quenched and then the MERFISH readout probes were sequentially hybridized and imaged in 2-3 colors.

To quench the fluorescent signal after each imaging round we used a toehold displacement strategy (as described previously^68^) in which unlabeled probes (IDT) complementary to the readout probes unhybridized the readout probes from the amplifier binding sites.

#### Sample preparation and hybridization

The cryosectioning, the hybridization of the encoding probe and the gel embedding and clearing of the sample were performed as described previously ^38^.

To fluorescently label the transcripts and amplify the signal, the preamplifier and amplifier probes were sequentially hybridized to the sample through the following protocol:

1. The 15 MERFISH preamplifiers and *Egr1* specific preamplifier were mixed together at a concentration of 30nM each in hybridization buffer (40% formamide, 10% dextran sulfate in 2xSSC) and hybridized to the sample for 30 minutes at 37 °C.
2. The sample was washed in 20% formamide in 2xSSC for 20 minutes at 37 °C.
3. The 15 MERFISH amplifiers and *Egr1* specific amplifier were mixed together at a concentration of 30nM each in hybridization buffer (40% formamide, 10% dextran sulfate in 2xSSC) and hybridized to the sample for 30 minutes at room temperature.
4. The sample was washed in 30% formamide in 2xSSC for 20 minutes at 37 °C.

Then to readout the MERFISH signal across combinations of OR transcripts, fluorescent readout probes were sequentially added to the sample using a fluidics device connected to an inverted epifluorescence microscope as described in ^67^. An identical labelling and imaging procedure was used for both the MOE and the OB.

#### MERFISH decoding of OR identity of sensory neurons in the MOE

The OR identity and volume occupied by each sensory neuron labelled was determined by first defining the best correlated groups of 3D pixels across each combination of 4 out of the set of 15 readout images corresponding to each MERFISH bit. These groups of pixels were determined by scanning ∼2.5 µm cubes across the epithelium and calculating all the Pearson correlation coefficients between the fluorescent signal across all the pairs of readout images. For these groups of pixels with high correlation (>0.6) we further applied a density-based clustering algorithm to group pixels corresponding to the same neuron and separate out distinct neurons. For this final filtered group of pixels, matching a particular set of 4 images, we found the OR identity based on the corresponding set of 4 readouts designed for the probes against the OR transcripts.

#### Imaging and aligning the OB sections

6 OB coronal sections were processed in each imaging experiment. Two pools of MERFISH probes (one targeting 500 ORs and the other targeting the remaining 600 ORs) were hybridized in an alternating manner across the sections.

To reduce imaging time, first a low magnification objective (10X) was used to image the nuclear signal (DAPI) across entire OB sections. This low-resolution overview of the sections allowed the high magnification (40X) MERFISH imaging to focus and tile only the glomeruli layer.

Approximately 160 coronal sections were imaged per animal, covering the bilateral OBs of 2 CD1 female mice (11 week) (Charles River) approximately uniformly in 16-18 µm increments. Each of the sections were aligned and corrected for sectioning distortions using the nuclear signal. First small subimages (of ∼ 100 μm in size) covering the glomeruli layer were individually rigidly aligned across adjacent sections by maximizing the correlation upon varying the angle and translation parameters. These served as matching features across the sections to be aligned. Using the consensus angle and translation, the adjacent slices were rigidly aligned. The remaining misalignment was corrected by gaussian interpolating the residual displacement of the matching features. Briefly for each point on the section, the additional distortion was calculated and corrected for from the residual displacement of the neighboring matching features which were weighed in using a gaussian term with a standard deviation ∼75 µm (similar to the glomeruli size). A custom graphics user interface was designed in Python and used to manually correct or exclude any features that were not correctly assigned via the automated procedure. By successively aligning each of the neighboring sections a 3D volume of the bulb can be reconstructed from the stack of ∼160 sections.

#### MERFISH decoding of OR identity of glomeruli in the OB

##### Glomeruli segmentation

The glomeruli within the glomerular layer of each coronal section of the olfactory bulb were segmented using Cellpose^46^a convolutional neural network-based algorithm designed primarily to segment cells/nuclei. This model was found, in general, to segment well roughly circular objects other than cells and nuclei. Briefly, the nuclear fluorescent signal (stained with DAPI) of the cells within the OB sections was inverted and rescaled to meet the dimension specifications of the pretrain Cellpose model. The nuclear pretrained model was used with low cutoff probability for the segmentation to ensure that the entire glomeruli layer is partitioned into glomeruli.

#### MERFISH staining into the olfactory bulb

Multiplexing was performed at a scale of 500-600 genes per section across two orthogonal sets of ORs. Glomeruli are typically 70-100 µm in size and cover a few successive sections. Therefore, by alternating between the two successive sets of probes used for hybridization across the serial sections, each glomerulus was interrogated across the entire 1,100 receptor repertoire.

#### Glomeruli decoding

Across the glomeruli layer, individual transcripts were identified using an algorithm for co-localizing 3D local maxima ^67^. Briefly, after localizing single molecules in 3D in each MERFISH readout image, we filter only the molecules co-localizing within ∼160 nm (across the x,y and z axes) in at least 4 out of 15 MERFISH readout images. For the molecules that colocalize across more than 4 MERFISH readout images we consider only the brightest 4. For each of these filtered molecules their OR identity is determined by matching their combination of 4 colocalizing MERFISH readouts to the unique combination of 4 readouts assigned during the MERFISH probe design for each OR gene. A brightness threshold is determined for each experiment to best separate the brightnesses of molecules mapping to the 500-600 ORs targeted vs the ORs not targeted during that experiment. This brightness threshold was applied to filter out the dim molecules contributing to noise.

For each segmented glomerulus we counted the number of transcripts of each OR and assigned it the identity of the most enriched OR if the number of identified transcripts was higher than 10 and the enrichment of the number of transcripts within the glomerulus was higher than 10 times over the other glomeruli within the section. The glomeruli identified in each section were mapped into the 3D volume of the OB reconstructed as described in the section above.

A few expected projections were not identified (i.e. projections of Olfr15 not detected in one of the bulbs). Overall, projections for ∼60% OR genes across the imaged OBs were first identified in this way. A potential source affecting the detection efficiency of OR glomerulus identity is the uneven distribution of the number of OR transcripts present in each glomerulus. Indeed, while for certain glomeruli we could reliably identify >50 OR transcripts per glomerular structure (Figure 3B,C), for other OR genes the corresponding glomeruli had less than 10 OR transcripts in each 2D coronal section, which was is below the limit of reliable detection. To increase the coverage we relaxed the identification criteria to more than 5 molecules instead of 10 if the same OR identity glomerulus was consistently identified within a similar position (within 500 µm) across multiple OBs.

#### Gene expression imputation in the MOE and OB

Single-cell sequencing data from cells isolated from the MOE of six C57BL/6J animals^47^ were used for calculating gene expression imputations in the MOE and OB. The previously published annotation^47^ was employed to exclude non-OSN cells and classify OSNs. For each OR/TAAR gene, the expression levels of each gene were averaged across their corresponding OSN populations. The average gene expression values for each OSN type were then spatially transferred to individual OSN positions identified by MERFISH across a pair of aligned reference MOE slices, capturing comprehensively the OR repertoire. Subsequently, spatially transferred gene expressions were averaged at each point on the MOE surface, considering all OSNs located within a 50 µm radius. A similar strategy was employed for OB surface mapping, where OSN-type averaged gene expressions were assigned to each OSN glomerular projection. Expression values were again spatially averaged within a 400 µm radius around each OB surface point. Finally, gene expression imputation values were normalized by dividing by the 99th percentile of the signal across each surface.

#### Predicting glomerular positions based on gene expression and imputation

To determine if differential gene expression across OSNs is predictive of the location of their OB projection, we modified the gene expression imputation approach described above to remove the contribution of the OSN type being evaluated. Specifically, for each OSN type, gene expression imputations were recalculated by transferring and averaging gene expression data from all other OSN types, excluding the selected OSN type. Specifically, we constructed gene expression imputations by transferring and averaging over the gene expression of all the other OSN projections excluding the OSN type selected. We then computed Pearson correlation coefficients between these modified imputations across the OB surface and the average gene expression profile of the selected OSN type (Figure S9A). Positions exhibiting the highest correlations within both the medial and lateral regions of the OB reference surface were identified as predicted projection locations based on gene expression.

#### Olfactory cue exposures

Adult (8-11 week) C57BL/6J (Jackson Laboratories) or CD1 mice (Charles River), including both females and males, were used in this study. Prior to exposures, all animals except mothers, were single housed for 2-7 days prior. CD1 mothers (2-5 days post birth) were single-housed into the same cage for 24h upon removing their pups from the home-cage. The cue was introduced in the home-cage of the animal to stimulate investigation of the foreign cue and minimize the introduction of other cues. After 30 minutes of exposure, in which the subject mice were allowed to freely interact, the animals were sacrificed. Then the MOE/OB was dissected (within 5-10 minutes) and freshly embedded in OCT (Tissue-Tek) and frozen in dry ice. The MOE was placed in an Eppendorf tube in PBS and degassed for 1 minute to remove air.

#### Acetophenone exposure

Following an odor exposure procedure published previously^31^, we spotted 10µL of undiluted acetophenone solution onto a piece of absorbing paper encased into a cassette and introduced it into the cage housing an 8-week C57BL/6J male. As in ^31^, we used 2 distinct concentrations of acetophenone in the spotted solution: undiluted 100% acetophenone and 1% acetophenone (diluted in DMSO), following the protocol in ^31^.

#### Cat bedding exposure

∼50 mL of soiled bedding was freshly sampled from the litter box of an adult female cat and introduced into the cage of the mouse (8-week C57BL/6J male) as described above. Bedding material was used expecting to contain a naturalistic mixture of olfactory odors excreted from the animal, including urine, feces, saliva, and fur.

#### Social cue exposure

A single C57BL/6J male or female mouse (8-11 weeks), or 4-5 pups (age postnatal day 2-5) were introduced into the home-cage of a CD1 mouse (male or female, 8 weeks). The animals were allowed to freely interact for 30 minutes after which the CD1 subject mice were sacrificed and their MOEs harvested as described above.

## Bibliography

1. Mori, K., and Sakano, H. (2021). Olfactory circuitry and behavioral decisions. Annu. Rev. Physiol. 83, 231–256.

2. Murthy, V.N. (2011). Olfactory maps in the brain. Annu. Rev. Neurosci. 34, 233–258.

3. Buck, L., and Axel, R. (1991). A novel multigene family may encode odorant receptors: a molecular basis for odor recognition. Cell 65, 175–187.

4. Chess, A., Simon, I., Cedar, H., and Axel, R. (1994). Allelic inactivation regulates olfactory receptor gene expression. Cell 78, 823–834.

5. Monahan, K., and Lomvardas, S. (2015). Monoallelic expression of olfactory receptors. Annu. Rev. Cell Dev. Biol. 31, 721–740.

6. Lomvardas, S., and Maniatis, T. (2016). Histone and DNA Modifications as Regulators of Neuronal Development and Function. Cold Spring Harb. Perspect. Biol. 8. 10.1101/cshperspect.a024208.

7. Monahan, K., Horta, A., and Lomvardas, S. (2019). LHX2- and LDB1-mediated trans interactions regulate olfactory receptor choice. Nature 565, 448–453.

8. Tan, L., Xie, X.S., and Lomvardas, S. (2025). Genomic snowflakes: how the uniqueness of DNA folding allows us to smell the chemical universe. Curr. Opin. Genet. Dev. 92, 102329.

9. Vassar, R., Chao, S., Sitcheran, R., Nuñez, J.M., Vosshall, L., and Axel, R. (1994). Topographic organization of sensory projections to the olfactory bulb. Cell 79, 981–991.

10. Ressler, K.J., Sullivan, S.L., and Buck, L.B. (1994). Information coding in the olfactory system: evidence for a stereotyped and highly organized epitope map in the olfactory bulb. Cell 79, 1245–1255.

11. Mombaerts, P., Wang, F., Dulac, C., Chao, S.K., Nemes, A., Mendelsohn, M., Edmondson, J., and Axel, R. (1996). Visualizing an olfactory sensory map. Cell 87, 675–686.

12. Ressler, K.J., Sullivan, S.L., and Buck, L.B. (1993). A zonal organization of odorant receptor gene expression in the olfactory epithelium. Cell 73, 597–609.

13. Vassar, R., Ngai, J., and Axel, R. (1993). Spatial segregation of odorant receptor expression in the mammalian olfactory epithelium. Cell 74, 309–318.

14. Zapiec, B., and Mombaerts, P. (2020). The zonal organization of odorant receptor gene choice in the main olfactory epithelium of the mouse. Cell Rep. 30, 4220–4234.e5.

15. Miyamichi, K., Serizawa, S., Kimura, H.M., and Sakano, H. (2005). Continuous and overlapping expression domains of odorant receptor genes in the olfactory epithelium determine the dorsal/ventral positioning of glomeruli in the olfactory bulb. J. Neurosci. 25, 3586–3592.

16. Tan, L., and Xie, X.S. (2018). A near-complete spatial map of olfactory receptors in the mouse main olfactory epithelium. Chem. Senses 43, 427–432.

17. Ruiz Tejada Segura, M.L., Abou Moussa, E., Garabello, E., Nakahara, T.S., Makhlouf, M., Mathew, L.S., Wang, L., Valle, F., Huang, S.S.Y., Mainland, J.D., et al. (2022). A 3D transcriptomics atlas of the mouse nose sheds light on the anatomical logic of smell. Cell Rep. 38, 110547.

18. Bashkirova, E.V., Klimpert, N., Monahan, K., Campbell, C.E., Osinski, J., Tan, L., Schieren, I., Pourmorady, A., Stecky, B., Barnea, G., et al. (2023). Opposing, spatially-determined epigenetic forces impose restrictions on stochastic olfactory receptor choice. Elife 12. 10.7554/eLife.87445.

19. Soucy, E.R., Albeanu, D.F., Fantana, A.L., Murthy, V.N., and Meister, M. (2009). Precision and diversity in an odor map on the olfactory bulb. Nat. Neurosci. 12, 210–220.

20. Zapiec, B., and Mombaerts, P. (2015). Multiplex assessment of the positions of odorant receptor-specific glomeruli in the mouse olfactory bulb by serial two-photon tomography. Proc. Natl. Acad. Sci. U. S. A. 112, E5873–82.

21. Pacifico, R., Dewan, A., Cawley, D., Guo, C., and Bozza, T. (2012). An olfactory subsystem that mediates high-sensitivity detection of volatile amines. Cell Rep. 2, 76–88.

22. Wang, I.-H., Murray, E., Andrews, G., Jiang, H.-C., Park, S.J., Donnard, E., Durán-Laforet, V., Bear, D.M., Faust, T.E., Garber, M., et al. (2022). Spatial transcriptomic reconstruction of the mouse olfactory glomerular map suggests principles of odor processing. Nat. Neurosci. 25, 484–492.

23. Zhu, K.W., Burton, S.D., Nagai, M.H., Silverman, J.D., de March, C.A., Wachowiak, M., and Matsunami, H. (2022). Decoding the olfactory map through targeted transcriptomics links murine olfactory receptors to glomeruli. Nat. Commun. 13, 5137.

24. Rubin, B.D., and Katz, L.C. (1999). Optical imaging of odorant representations in the mammalian olfactory bulb. Neuron 23, 499–511.

25. Belluscio, L., and Katz, L.C. (2001). Symmetry, stereotypy, and topography of odorant representations in mouse olfactory bulbs. J. Neurosci. 21, 2113–2122.

26. Meister, M., and Bonhoeffer, T. (2001). Tuning and topography in an odor map on the rat olfactory bulb. J. Neurosci. 21, 1351–1360.

27. Wachowiak, M., Denk, W., and Friedrich, R.W. (2004). Functional organization of sensory input to the olfactory bulb glomerulus analyzed by two-photon calcium imaging. Proc. Natl. Acad. Sci. U. S. A. 101, 9097–9102.

28. Mori, K., Takahashi, Y.K., Igarashi, K.M., and Yamaguchi, M. (2006). Maps of odorant molecular features in the Mammalian olfactory bulb. Physiol. Rev. 86, 409–433.

29. Ma, L., Qiu, Q., Gradwohl, S., Scott, A., Yu, E.Q., Alexander, R., Wiegraebe, W., and Yu, C.R. (2012). Distributed representation of chemical features and tunotopic organization of glomeruli in the mouse olfactory bulb. Proc. Natl. Acad. Sci. U. S. A. 109, 5481–5486.

30. Peterlin, Z., Firestein, S., and Rogers, M.E. (2014). The state of the art of odorant receptor deorphanization: a report from the orphanage. J. Gen. Physiol. 143, 527–542.

31. Jiang, Y., Gong, N.N., Hu, X.S., Ni, M.J., Pasi, R., and Matsunami, H. (2015). Molecular profiling of activated olfactory neurons identifies odorant receptors for odors in vivo. Preprint, 10.1038/nn.4104 10.1038/nn.4104.

32. von der Weid, B., Rossier, D., Lindup, M., Tuberosa, J., Widmer, A., Col, J.D., Kan, C., Carleton, A., and Rodriguez, I. (2015). Large-scale transcriptional profiling of chemosensory neurons identifies receptor-ligand pairs in vivo. Nat. Neurosci. 18, 1455.

33. Vihani, A., Hu, X.S., Gundala, S., Koyama, S., Block, E., and Matsunami, H. (2020). Semiochemical responsive olfactory sensory neurons are sexually dimorphic and plastic. Elife 9. 10.7554/eLife.54501.

34. Hu, X.S., Ikegami, K., Vihani, A., Zhu, K.W., Zapata, M., de March, C.A., Do, M., Vaidya, N., Kucera, G., Bock, C., et al. (2020). Concentration-dependent recruitment of mammalian odorant receptors. eNeuro 7, ENEURO.0103-19.2019.

35. Chen, K.H., Boettiger, A.N., Moffitt, J.R., Wang, S., and Zhuang, X. (2015). RNA imaging. Spatially resolved, highly multiplexed RNA profiling in single cells. Science 348, aaa6090.

36. Isogai, Y., Si, S., Pont-Lezica, L., Tan, T., Kapoor, V., Murthy, V.N., and Dulac, C. (2011). Molecular organization of vomeronasal chemoreception. Nature 478, 241–245.

37. van der Linden, C., Jakob, S., Gupta, P., Dulac, C., and Santoro, S.W. (2018). Sex separation induces differences in the olfactory sensory receptor repertoires of male and female mice. Nat. Commun. 9, 5081.

38. Moffitt, J.R., Bambah-Mukku, D., Eichhorn, S.W., Vaughn, E., Shekhar, K., Perez, J.D., Rubinstein, N.D., Hao, J., Regev, A., Dulac, C., et al. (2018). Molecular, spatial, and functional single-cell profiling of the hypothalamic preoptic region. Science 362, eaau5324.

39. Xia, C., Fan, J., Emanuel, G., Hao, J., and Zhuang, X. (2019). Spatial transcriptome profiling by MERFISH reveals subcellular RNA compartmentalization and cell cycle-dependent gene expression. Proc. Natl. Acad. Sci. U. S. A. 116, 19490–19499.

40. Ibarra-Soria, X., Levitin, M.O., Saraiva, L.R., and Logan, D.W. (2014). The olfactory transcriptomes of mice. PLoS Genet. 10, e1004593.

41. Goh, J.J.L., Chou, N., Seow, W.Y., Ha, N., Cheng, C.P.P., Chang, Y.-C., Zhao, Z.W., and Chen, K.H. (2020). Highly specific multiplexed RNA imaging in tissues with split-FISH. Nat. Methods 17, 689–693.

42. Xia, C., Babcock, H.P., Moffitt, J.R., and Zhuang, X. (2019). Multiplexed detection of RNA using MERFISH and branched DNA amplification. Sci. Rep. 9, 7721.

43. Vassalli, A., Rothman, A., Feinstein, P., Zapotocky, M., and Mombaerts, P. (2002). Minigenes impart odorant receptor-specific axon guidance in the olfactory bulb. Neuron 35, 681–696.

44. Markenscoff-Papadimitriou, E., Allen, W.E., Colquitt, B.M., Goh, T., Murphy, K.K., Monahan, K., Mosley, C.P., Ahituv, N., and Lomvardas, S. (2014). Enhancer interaction networks as a means for singular olfactory receptor expression. Cell 159, 543–557.

45. Pyrski, M., Xu, Z., Walters, E., Gilbert, D.J., Jenkins, N.A., Copeland, N.G., and Margolis, F.L. (2001). The OMP-lacZ transgene mimics the unusual expression pattern of OR-Z6, a new odorant receptor gene on mouse chromosome 6: implication for locus-dependent gene expression. J. Neurosci. 21, 4637–4648.

46. Stringer, C., Wang, T., Michaelos, M., and Pachitariu, M. (2021). Cellpose: a generalist algorithm for cellular segmentation. Nat. Methods 18, 100–106.

47. Tsukahara, T., Brann, D.H., Pashkovski, S.L., Guitchounts, G., Bozza, T., and Datta, S.R. (2021). A transcriptional rheostat couples past activity to future sensory responses. Cell 184, 6326–6343.e32.

48. Coleman, J.H., Lin, B., Louie, J.D., Peterson, J., Lane, R.P., and Schwob, J.E. (2019). Spatial determination of neuronal diversification in the olfactory epithelium. J. Neurosci. 39, 814–832.

49. Mountoufaris, G., Chen, W.V., Hirabayashi, Y., O’Keeffe, S., Chevee, M., Nwakeze, C.L., Polleux, F., and Maniatis, T. (2017). Multicluster Pcdh diversity is required for mouse olfactory neural circuit assembly. Science 356, 411–414.

50. Gene Ontology Consortium (2010). The Gene Ontology in 2010: extensions and refinements. Nucleic Acids Res. 38, D331–5.

51. Sakano, H. (2020). Developmental regulation of olfactory circuit formation in mice. Dev. Growth Differ. 62, 199–213.

52. Assens, A., Dal Col, J.A., Njoku, A., Dietschi, Q., Kan, C., Feinstein, P., Carleton, A., and Rodriguez, I. (2016). Alteration of Nrp1 signaling at different stages of olfactory neuron maturation promotes glomerular shifts along distinct axes in the olfactory bulb. Development 143, 3817–3825.

53. Cho, J.H., Kam, J.W.K., and Cloutier, J.-F. (2012). Slits and Robo-2 regulate the coalescence of subsets of olfactory sensory neuron axons within the ventral region of the olfactory bulb. Dev. Biol. 371, 269–279.

54. McIntyre, J.C., Titlow, W.B., and McClintock, T.S. (2010). Axon growth and guidance genes identify nascent, immature, and mature olfactory sensory neurons: Developmental Regulation of Axon Guidance Genes. J. Neurosci. Res. 88, 3243–3256.

55. Mosca, T.J., and Luo, L. (2014). Synaptic organization of the Drosophila antennal lobe and its regulation by the Teneurins. Elife 3, e03726.

56. Liberles, S.D., and Buck, L.B. (2006). A second class of chemosensory receptors in the olfactory epithelium. Nature 442, 645–650.

57. Dewan, A., Pacifico, R., Zhan, R., Rinberg, D., and Bozza, T. (2013). Non-redundant coding of aversive odours in the main olfactory pathway. Nature 497, 486–489.

58. Li, Q., Korzan, W.J., Ferrero, D.M., Chang, R.B., Roy, D.S., Buchi, M., Lemon, J.K., Kaur, A.W., Stowers, L., Fendt, M., et al. (2013). Synchronous evolution of an odor biosynthesis pathway and behavioral response. Curr. Biol. 23, 11–20.

59. Hanchate, N.K., Kondoh, K., Lu, Z., Kuang, D., Ye, X., Qiu, X., Pachter, L., Trapnell, C., and Buck, L.B. (2015). Single-cell transcriptomics reveals receptor transformations during olfactory neurogenesis. Science 350, 1251–1255.

60. Saraiva, L.R., Ibarra-Soria, X., Khan, M., Omura, M., Scialdone, A., Mombaerts, P., Marioni, J.C., and Logan, D.W. (2015). Hierarchical deconstruction of mouse olfactory sensory neurons: from whole mucosa to single-cell RNA-seq. Sci. Rep. 5, 18178.

61. Schaefer, M., Yamazaki, K., Osada, K., Restrepo, D., and Beauchamp, G. (2002). Olfactory fingerprints for major histocompatibility complex-determined body odors II: Relationship among odor maps, genetics, odor composition, and behavior. J. Neurosci. 22, 9513–9521.

62. Matsuo, T., Hattori, T., Asaba, A., Inoue, N., Kanomata, N., Kikusui, T., Kobayakawa, R., and Kobayakawa, K. (2015). Genetic dissection of pheromone processing reveals main olfactory system-mediated social behaviors in mice. Proc. Natl. Acad. Sci. U. S. A. 112, E311–20.

63. Takahashi, L.K. (2014). Olfactory systems and neural circuits that modulate predator odor fear. Front. Behav. Neurosci. 8, 72.

64. Aoki, M., Gamayun, I., Wyatt, A., Grünewald, R., Simon-Thomas, M., Philipp, S.E., Hummel, O., Wagenpfeil, S., Kattler, K., Gasparoni, G., et al. (2021). Prolactin-sensitive olfactory sensory neurons regulate male preference in female mice by modulating responses to chemosensory cues. Sci. Adv. 7, eabg4074.

65. Lee, D., Kume, M., and Holy, T.E. (2019). Sensory coding mechanisms revealed by optical tagging of physiologically defined neuronal types. Science 366, 1384–1389.

66. Emanuel, G., seichhorn, Babcock, H., leonardosepulveda, and timblosser (2020). ZhuangLab/MERlin: MERlin v0.1.6 (Zenodo) 10.5281/ZENODO.3758540.

67. Su, J.-H., Zheng, P., Kinrot, S.S., Bintu, B., and Zhuang, X. (2020). Genome-scale imaging of the 3D organization and transcriptional activity of chromatin. Cell 182, 1641–1659.e26.

68. Bintu, B., Mateo, L.J., Su, J.-H., Sinnott-Armstrong, N.A., Parker, M., Kinrot, S., Yamaya, K., Boettiger, A.N., and Zhuang, X. (2018). Super-resolution chromatin tracing reveals domains and cooperative interactions in single cells. Science 362. 10.1126/science.aau1783.

